# Competence remodels the pneumococcal cell wall providing resistance to fratricide and surface exposing key virulence factors

**DOI:** 10.1101/2022.08.03.502730

**Authors:** Vikrant Minhas, Arnau Domenech, Dimitra Synefiaridou, Daniel Straume, Max Brendel, Gonzalo Cebrero, Xue Liu, Charlotte Costa, Mara Baldry, Jean-Claude Sirard, Camilo Perez, Nicolas Gisch, Sven Hammerschmidt, Leiv Sigve Håvarstein, Jan-Willem Veening

**Author notes:** Correspondence to or; Twitter: @JWVeening. These authors contributed equally (Vikrant Minhas, Arnau Domenech, Dimitra Synefiaridou).

## Abstract

Competence development in the human pathogen *Streptococcus pneumoniae* controls several features such as genetic transformation, biofilm formation and virulence. Competent bacteria produce so called ‘fratricins’ such as CbpD, that kill non-competent siblings by cleaving peptidoglycan (PGN). CbpD is a choline-binding protein (CBP) that binds to phosphorylcholine residues found on wall- and lipoteichoic acids (WTA and LTA) that together with PGN are major constituents of the pneumococcal cell wall. Competent pneumococci are protected against fratricide by producing the immunity protein ComM. How competence and fratricide contribute to virulence is unknown. Here, using a genome-wide CRISPRi-seq screen, we show that genes involved in teichoic acid biosynthesis are essential during competence. We demonstrate that LytR is the major enzyme mediating the final step in WTA formation, and that, together with ComM, is essential for immunity against CbpD. Importantly, we show that key virulence factors PspA and PspC become more surface-exposed at midcell during competence, in a CbpD-dependent manner. Together, our work supports a model in which activation of competence is crucial for host adherence by increased surface exposure of its various CBPs.

## Introduction

*Streptococcus pneumoniae* (the pneumococcus) is a member of the commensal microbiota of the human nasopharynx. However, it is a major public health problem because it can cause severe life-threatening infections such as sepsis, pneumonia and meningitis (Wahl et al., 2018). The pneumococcus is a naturally transformable bacterium that can take up and assimilate exogenous DNA (Chewapreecha et al., 2014; Dawson and Sia, 1931; Johnston et al., 2014; Veening and Blokesch, 2017). This phenomenon is an important mechanism of genome plasticity and is largely responsible for the acquisition and spread of antibiotic resistance as well as virulence factors such as the capsule (Croucher et al., 2011; Wyres et al., 2013). The transformation process in the pneumococcus requires the induction of a physiological state named competence, which involves about 10% of the pneumococcal genome (Claverys et al., 2009; Slager et al., 2019).

Competence is induced by a classical two-component quorum sensing system in which the *comC*-encoded competence-stimulating peptide (CSP) is cleaved and exported by the membrane transporter ComAB to the extracellular space (Håvarstein et al., 1995; Hui and Morrison, 1991). Upon a certain threshold of CSP accumulation, it stimulates the autophosphorylation of the membrane-bound histidine-kinase ComD, which subsequently activates its cognate response regulator ComE (Håvarstein et al., 1996; Martin et al., 2013; Pestova et al., 1996). Phosphorylated ComE induces a positive feedback to activate the early *com* genes including its own operon. One of the genes regulated by ComE, *comX*, encodes a sigma factor that activates the late *com* genes required for DNA repair, DNA uptake, and transformation (Lee and Morrison, 1999; Martin et al., 2013). Although competence is mostly associated with DNA transformation, only 22 of the genes induced during competence are related to transformation (Dagkessamanskaia et al., 2004; Peterson et al., 2004). Other competence genes are involved in biofilm formation (Aggarwal et al., 2018), bacteriocin production (Kjos et al., 2016; Wholey et al., 2016) and siblings’ fratricide (Steinmoen et al., 2002).

As pneumococci do not discriminate between homologous and foreign DNA, it is believed that fratricide serves to increase and facilitate the exchange of DNA between pneumococci. Thus, competent pneumococci lyse and subsequently release nutrients and DNA from a subfraction of the population that does not become competent (Steinmoen et al., 2002). Three choline-binding proteins (CBPs) constitute the lysis mechanism: CbpD, LytA, and LytC. CbpD is the main driver of fratricide as the process cannot commence in its absence (Eldholm et al., 2009; Kausmally et al., 2005). CBPs are non-covalently bound to the phosphorylcholine (*P*Cho) of the teichoic acid (Galán-Bartual et al., 2015). CbpD, LytA and LytC bind to the *P*Cho residues in the wall teichoic acids (WTA) that are covalently attached to the peptidoglycan (PGN) or to the lipid-anchored lipoteichoic acids (LTA) (Kausmally et al., 2005; Swiatlo et al., 2002; Weidenmaier and Peschel, 2008). As most non-pneumococci do not contain choline in their cell walls, lysis by fratricins enhances the probability for accessing homologous DNA (Johnsborg et al., 2008). The pneumococcus is special in its teichoic acid biosynthesis pathway, as the same substrate is used for LTA and WTA (Figure 1A) (Denapaite et al., 2012). It was recently shown that the lipoteichoic acid ligase TacL is responsible for producing LTA by transferring the polymeric TA chain from an undecaprenyl-diphosphate (C55-PP) linked precursor onto the glycolipid anchor (Flores-Kim et al., 2019; Heß et al., 2017). The three putative enzymes of the LCP-protein family Cps2A, Psr, and LytR are predicted to anchor the polymeric TA chain from the same precursor to the PGN to form WTA, but it remains unclear which is the dominant route and how the levels of WTA and LTA are controlled (Figure 1B) (Denapaite et al., 2012; Eberhardt et al., 2012; Kawai et al., 2011). In addition, it is assumed that most pneumococcal strains – as with other Gram-positive bacteria - anchor their exopolysaccharide capsule on the same residue within the PGN as WTA, that would make a cross-talk between these two processes likely (Eberhardt et al., 2012; Stefanović et al., 2021). By contrast, a different attachment position for the capsule in *S. pneumoniae* has been suggested recently (Larson and Yother, 2017) but final structural proof on the molecular level, e.g. by two-dimensional NMR experiments, still remains to be demonstrated.

**Figure 1.**
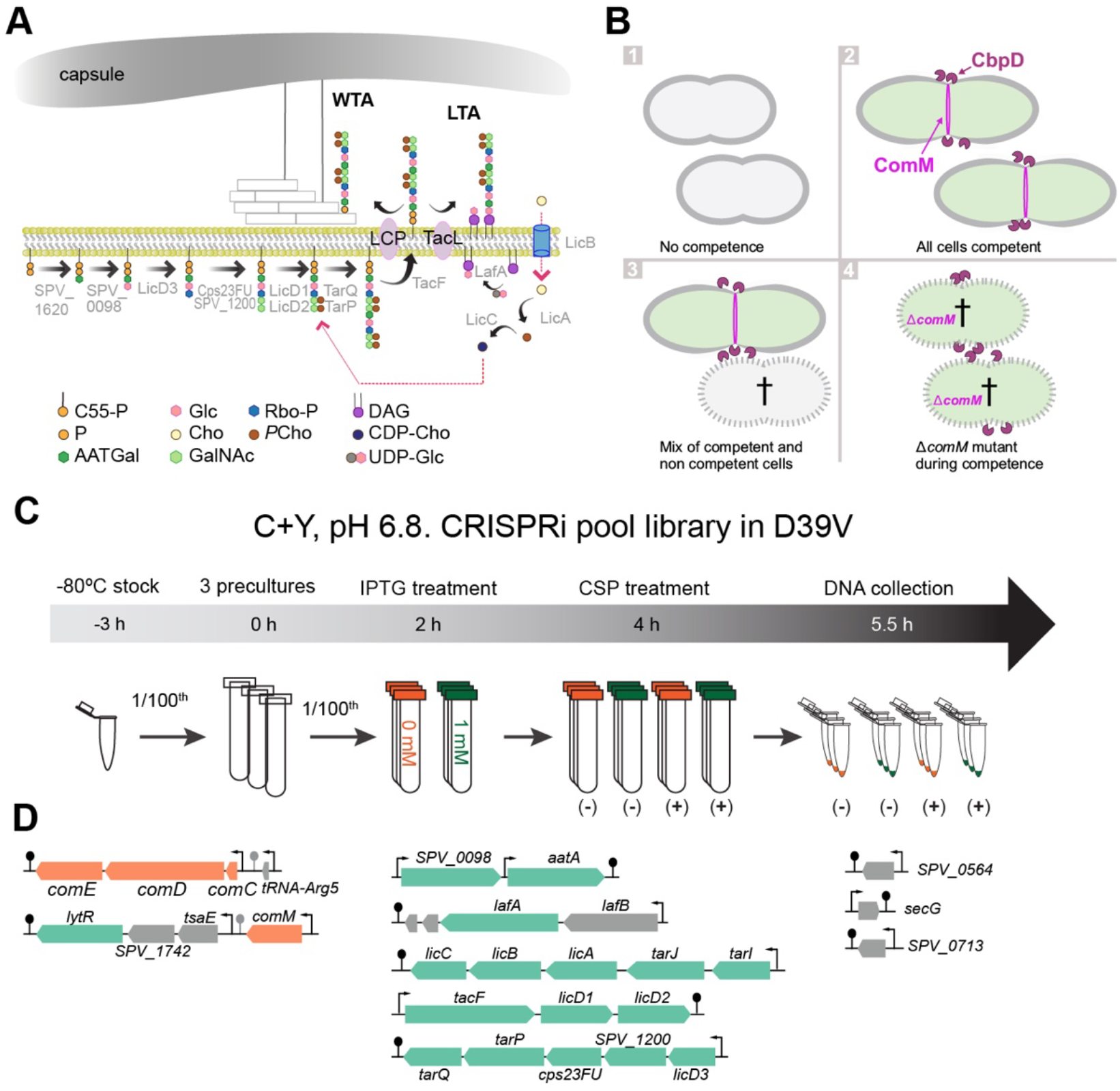
**A)** Proposed teichoic acid biosynthesis pathway in *S. pneumoniae*. Abbreviations: LCP (LytR-Cps2A-Psr family), C55-PP (Undecaprenyl diphosphate), P (Phosphate residue), Glc (Glucose), Rbo-P (ribitol 5-phosphate), DAG (Diacylglycerol), AATGal (2-acetamido-4-amino-2,4,6-trideoxygalactose), Cho (Choline), *P*Cho (phosphorylcholine), CDP-Cho (cytidine diphosphate-choline), GalNac (N-acetylgalactosamine), UDP-Glc (uridine diphosphate glucose). Note that the capsule and the WTA potentially compete for the same anchoring position on the peptidoglycan (O-6 of N-acetylmuramic acid (MurNAc)). **B)** Pneumococcal fratricide. 1) under non-permissive conditions for competence, D39V cells do not become competent, thus, the fratricin CbpD and the immunity protein ComM are not expressed. 2) Under permissive conditions, all cells become competent, producing CbpD but also ComM leading to cell elongation (Bergé et al., 2017), thus all cells are protected and there is no fratricide. 3) In conditions where only a subpopulation of cells become competent, non-competent pneumococci lyse by action of CbpD and subsequently release DNA. 4) Deletion of *comM* results in autolysis when competence is activated via CbpD production. **C)** Workflow to detect essential genes during competence. Three independent precultures of the CRISPRi pooled library (Liu et al., 2021) were grown until OD 0.1 in C+Y pH 6.8 to avoid spontaneous competence. Cells were then diluted 100 times in C+Y supplemented with 0 or 1 mM of IPTG. When cells reached OD_595 nm_ 0.1 again, 100 ng/ml CSP_1_ was added to the indicated conditions and cells were grown for two more generations (OD_595 nm_ ~ 0.4). Cells were collected, the DNA was isolated, and the library was prepared for Illumina sequencing (de Bakker et al., 2022). **D)** Operons with fitness cost during competence. In orange, competence-related genes; in green, genes involved in teichoic acid biosynthesis.

CbpD, which is only expressed during competence, consists of four domains: the N-terminal cysteine/histidine-dependent amidohydrolase-peptidase (CHAP) domain that most likely cleaves peptide bonds of the PGN chain (Eldholm et al., 2009, 2010); the two src-homology 3b (SH3b) domains that recognize and bind to PGN; and the C-terminal end consisting of four repeating choline binding sequences that direct CbpD to the septal region of the pneumococcal cell (Eldholm et al., 2010). The initial cell wall damage by CbpD most likely facilitates the accessibility of LytA and LytC, enhancing the fratricide process. Interestingly, several key virulence factors such as CbpG (cleaves host extracellular matrix (Mann et al., 2006)), pneumococcal surface protein A (PspA - inhibits opsonisation and binding to host lactate dehydrogenase (Park et al., 2021)) and PspC (aka CbpA - involved in adherence and complement evasion (Elm et al., 2004; Iovino et al., 2017; Rosenow et al., 1997; Zhang et al., 2000)) are also CBPs (Hakenbeck et al., 2009; Kadioglu et al., 2008). Although not understood, a *cbpD* mutant is attenuated in colonization (Gosink et al., 2000).

To avoid committing suicide during fratricide, competent cells produce ComM, an immunity protein (Håvarstein et al., 2006). During competence, ComM (ComE-dependent) is produced earlier (~5 min) than CbpD (ComX-dependent). ComM is an integral transmembrane protein, of yet unknown structure, whose role in providing immunity still remains to be elucidated. Some clues come from two studies that showed that cells become elongated during competence in a ComM-dependent manner, suggesting that ComM regulates septal PGN synthesis (Bergé et al., 2017; Straume et al., 2017a) (Figure 1B).

In addition, competence is also activated when cells are adhering to human epithelial cells (Aprianto et al., 2016), and during invasive disease as shown in several mouse and zebrafish models (Lin et al., 2020; Schmidt et al., 2019)(Jim et al., 2022). Indeed, competence development is a key pathogenic factor in pneumococcal meningitis (Jim et al., 2022; Schmidt et al., 2019). However, it is unclear how competence and fratricide play a role in pneumococcal virulence.

Here, we found that teichoic acid biosynthesis is essential during competence. We demonstrate that ComM provides immunity against fratricins in concert with LytR, and that LytR is a key enzyme required for WTA anchoring in the pneumococcus. Transcriptional repression of any of the genes involved in WTA assembly will lead to increased susceptibility to CbpD and fratricide. Strikingly, we find that PspA and PspC are localised to the septum and are more readily detected by antibodies when bacteria are competent, with this phenomenon being dependent on CbpD. In line with CbpD-dependent surface exposure of CBPs, pneumococci lacking CbpD show reduced adherence to human nasopharyngeal epithelial cells. The data presented here suggest a model in which competence activation during infection is crucial to expose important virulence factors to the outside of the pneumococcal cell to ensure better adherence to host cells.

## Results

### Teichoic acid biosynthesis is essential during pneumococcal competence as identified by CRISPRi-seq

While it has been shown that the competence-induced ComM protein is required for immunity against fratricins (Håvarstein et al., 2006), the molecular mechanism underlying immunity remains unclear. To investigate this, we performed a genome-wide CRISPRi-seq screen (Figure 1C-D) (de Bakker et al., 2022). We screened a pooled library of pneumococcal strains harbouring an inducible dCas9 and a constitutively expressed sgRNA, targeting in total 1498 operons in the genome of strain D39V (Liu et al., 2021). The rationale behind this screen being that bacteria undergoing transcriptional downregulation of genes important for immunity against fratricins will be outcompeted or lysed by fratricins produced by strains in which the sgRNA targets a neutral gene (Figure 1B). The pooled library was grown under competence-permissive and non-permissive conditions by the addition of synthetic CSP_1_. To confirm that observed fitness costs were due to competence induction and not because of the essentiality of the targeted operon, four different conditions were tested (Figure S1A): (I) control (C+Y, pH 6.8; non-permissive for natural competence development), (II) library induction in absence of competence (+ IPTG), (III) competence induction (+ CSP_1_) and (IV) both library and competence induction (+ IPTG and CSP_1_). After Illumina sequencing, the fitness of targets and quality of the replicates were then evaluated and the fold change of the abundance of sgRNAs between the four groups was analysed (de Bakker et al., 2022)(Figure S1B-C).

Fourteen sgRNAs targeting ten operons were significantly less abundant in condition IV, suggesting that they are essential or become more essential during competence (Figures 1D and S2A, and Table S1). As expected, *comCDE* and *comM* were among the top hits, demonstrating that when cells are unable to activate competence or ComM, they are rapidly outcompeted or lysed by competent (and CbpD) producing competitors. Strikingly, most of the other hits were operons related to teichoic acid (TA) synthesis (Figure 1D).

Next, we examined in more detail the identified sgRNAs targeting genes involved in both competence and TA synthesis (Figure S2). All but two operons related to TA synthesis (*SPV_1620-aatB*, and *tacL*) were underrepresented (Figure S2A). The first operon was less present when competence and the library were induced, although the differences were not significant; however, the sgRNA targeting *tacL*, which transfers TA precursor chains onto the glycolipid membrane anchor (forming LTA) (Flores-Kim et al., 2019; Heß et al., 2017), did not show any fitness cost during competence (Figure S2B). The three remaining sgRNAs targeted *secG* and two hypothetical proteins: *SPV_0713* and *SPV_0564*.

### Teichoic acid biosynthesis does not regulate competence development

One hypothesis that could explain why strains carrying a sgRNA targeting a TA gene are under-represented in the CRISPRi-seq screen, is that competence is not activated and hence ComM is not expressed rendering cells susceptible to fratricide. To test whether TAs are required for competence activation, we generated non-polar depletion strains for each individual gene belonging to the 10 operons (Table S2). The deletions containing an erythromycin resistance cassette were transformed in the corresponding strain harbouring an IPTG-inducible copy of the gene (P*_lac_* promoter) integrated at the non-essential ZIP locus, in order to control the expression of the genes (e.g. *comM* ^-/Plac-^) (Figure S3). High-content microscopy of the depletion strains showed different phenotypes depending on the targeted gene, as shown before (Figure S4) (Liu et al., 2017). We used a competence-specific induced *ssbB* promoter fused to firefly luciferase (*P_ssbB_-luc*, (Slager et al., 2014) to quantify competence activation. Cells were grown in presence of different concentrations of IPTG to deplete or overproduce the TA gene products (0, 0.005 mM, 0.01 mM, 0.1 mM and 1 mM), and luciferase activity as well as optical density were regularly measured (Figure S5). As expected, in absence of *comCDE*, competence was not triggered. However, transcriptional repression of all tested TA-related genes, except *tacL*, did not affect natural competence development under conditions where cell growth was not compromised (Figure S5). Interestingly, although the absence of TacL inhibited natural competence, cells remained protected from fratricide when induced by externally added CSP (Figure S6), explaining why *tacL* depletion did not show any fitness cost in the CRISPRi experiment (Figure S2).

### Teichoic acid biosynthesis is required to prevent cell lysis by fratricide

As TAs are not required for competence induction (Figure S6), another hypothesis is that TAs provide resistance to fratricins during competence. In this case, depletion of TA-related genes should result in autolysis by fratricides when competence is triggered. To test this, we used the SYTOX™ Green Dead Cell Stain (ThermoFisher scientific) to evaluate cell lysis in presence and absence of CSP_1_-induced competence. A *comM* mutant was used as a control strain that should show lysis upon competence induction. Indeed, as shown in Figure S6, rapid lysis after CSP_1_ addition was observed for the *comM* mutant, as well as many of the TA-related genes (e.g. *tarP, tacF, tarI, licA, licD3*). This data suggests that, besides ComM, TAs are essential to maintain the integrity of the cells during ‘attack’ of competence-induced fratricins (i.e. CbpD, LytA). Indeed, it was previously shown that cells depleted for TAs become more susceptible to autolysis by LytA (Bonnet et al., 2018).

### lytR *is in a transcriptional unit with* comM *and is upregulated during competence by transcriptional read-through*

As both CbpD and LytA bind to the choline units on TAs, we focused on the external part of the pathway: the anchoring of TAs into cell wall (WTA) and membrane (LTA), respectively (Figure 2A). TacL is the only known protein in *S. pneumoniae* responsible for anchoring TA chains to a glycolipid, while LCP family proteins (i.e. LytR, Cps2A and Psr) are the putative proteins anchoring such chains to the cell wall thereby producing WTA. The fact that *cps2A* and *psr* were not found in the CRISPRi-seq screen (Figure S2), and that *lytR* forms part of the same operon as *comM* (Figure 2B), suggests a key role for LytR in attaching TAs into the cell wall and subsequent protection from fratricins.

**Figure 2.**
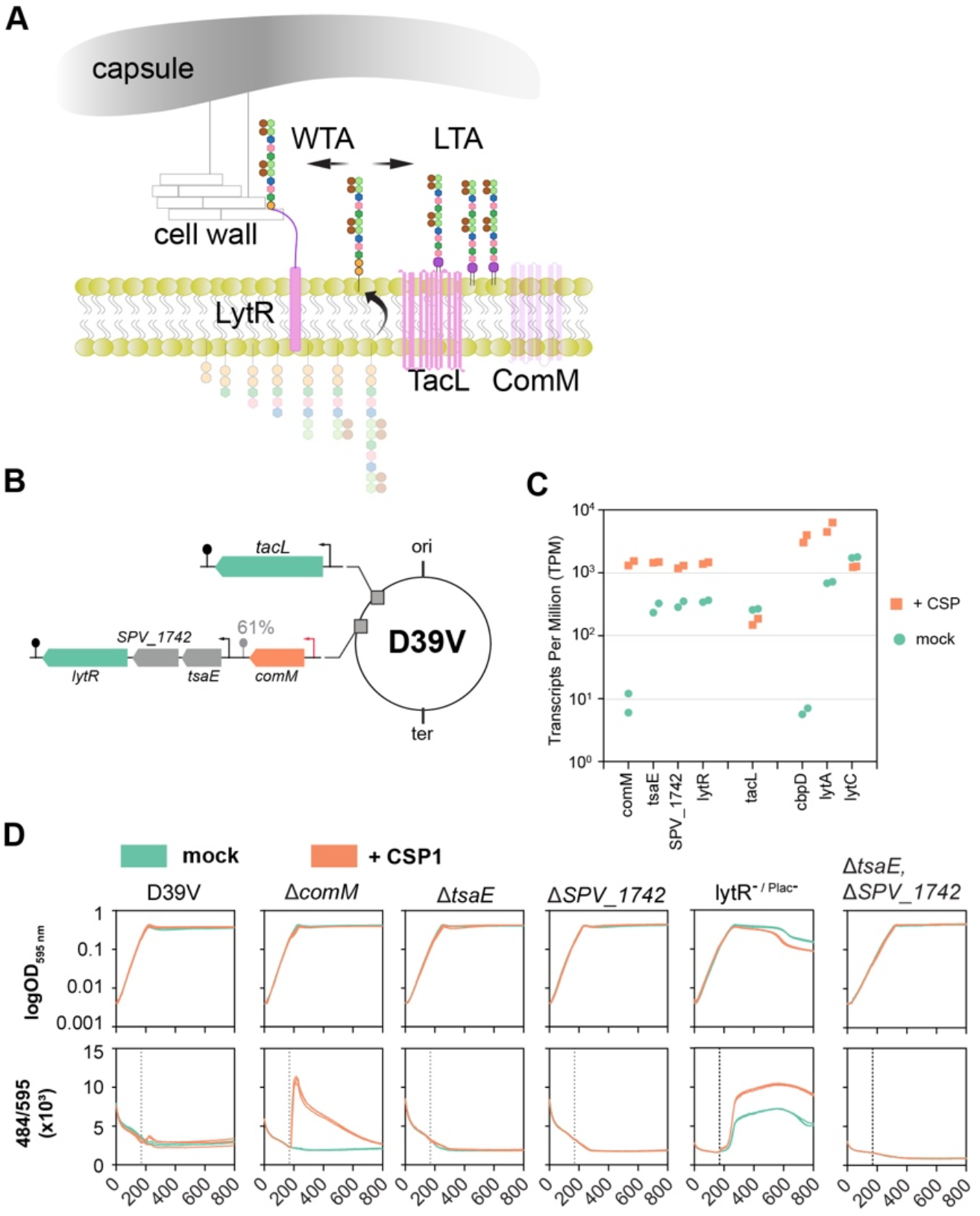
Role of the *comM-tsaE-SPV_1742-lytR* and *tacL* operons. **A)** In *S. pneumoniae* the same TA chains can be anchored to the membrane (by TacL) or to the cell wall with LytR as key enzyme. **B)** Schematic representation of the *tacL, comM* and *lytR* operons. Note the ComE-activated promoter (orange arrow) upstream of *comM* and the imperfect transcriptional terminator downstream of *comM* (Slager et al., 2018). **C)** Expression levels in competence induced and non-induced cells. RNA-seq data from (Aprianto et al., 2018; Slager et al., 2019). Duplicates for each condition are shown. **D)** Cell lysis evaluation. Individual strains were grown in C+Y pH 6.8 to avoid natural competence development in presence of SYTOX™ Green Dead Cell Stain dye. When cell cultures reached OD_595 nm_ 0.1 (~ after 170 min), 100 ng/ml of CSP_1_ was added to induce competence. Three biological replicates per condition are shown. As LytR is essential, the depletion strain (lytR ^- / Plac-^) was used in absence of IPTG, to deplete for LytR.

While *tacL* is in a single gene operon, the *comM-tsaE-spv_1742-lytR* operon shows an interesting complexity (Slager et al., 2018). Transcription of *comM* is under competence control by ComE, and its imperfect terminator is approximately 61% efficient (Figure 2B). The three downstream genes are regulated by an internal transcription start site (TSS) with basal expression. However, when competence is activated, the expression of all three genes is highly increased due to inefficient transcriptional termination by the *comM* terminator (Figure 2C; (Aprianto et al., 2018; Slager et al., 2019). We also note strong activation of *cbpD* transcription and to a lesser extent *lytA* during competence induction, while *tacL* and *lytC* transcription is not changed (Figure 2C). TsaE is a predicted tRNA threonylcarbamoyladenosine biosynthesis protein and SPV_1742 is a predicted acetyltransferase (Slager et al., 2018). Individual deletion of *tsaE* or *spv_1742* did not affect cell integrity during competence (Figures 2D and S5). To further exclude their role in fratricide, we constructed a double *ΔtsaE ΔSPV_1742* deletion mutant. As shown in Figure 2D, the *ΔtsaE ΔSPV_1742* double mutant was not susceptible to lysis during competence, suggesting they do not play a role in TA synthesis.

### LytR is required for ComM-mediated immunity to CbpD

While *tsaE* and *spv_1742* did not play a role in immunity to fratricins, we were unable to construct a *lytR* deletion strain in the D39V genetic background, suggesting it is an essential gene under our experimental conditions. We note that pneumococcal *lytR* mutants have been successfully constructed before, although serious growth impairments were noted (Johnsborg and Håvarstein, 2009; Ye et al., 2018). Nevertheless, we were able to construct a complementation strain in which *lytR* is expressed from an IPTG-inducible promoter at the ZIP locus (Keller et al., 2019) while deleted from its native locus (Figure S3). To evaluate how LytR protects cells from competence-related lysis, we performed three complementary approaches: evaluation of cell lysis in the deletion of the fratricins (Figure 3A), overexpression of *cbpD* (Figure 3B) and addition of exogenous CbpD (Figure 3C). In all three approaches, absence of competence or *comM* depletion resulted in a rapid cell lysis when CbpD was present (either induced or added exogenously to the medium). Importantly, depletion of *lytR* strongly increased susceptibility to CbpD (Figure 3C) in line with a previous report (Johnsborg and Håvarstein, 2009). Deletion of both *cbpD* and *lytA* in lytR^-/Plac-^ cells does not protect them from lysis, confirming the essential role of LytR, not only during competence (Figure 3A). Interestingly, LytR overexpression in the presence of basal levels of ComM showed some level of protection against CbpD-mediated lysis, which was more evident when both ComM and LytR were overexpressed (Figure 3). Indeed, overproduction of both ComM and LytR provided better protection from CbpD-induced lysis than overproduction of ComM alone (Figure 3C). In contrast, native expression and overexpression of LytR in the absence of ComM (*comM* depletion), showed the same levels of lysis as a *comM* mutant, suggesting that both proteins are required to protect from fratricide.

**Figure 3.**
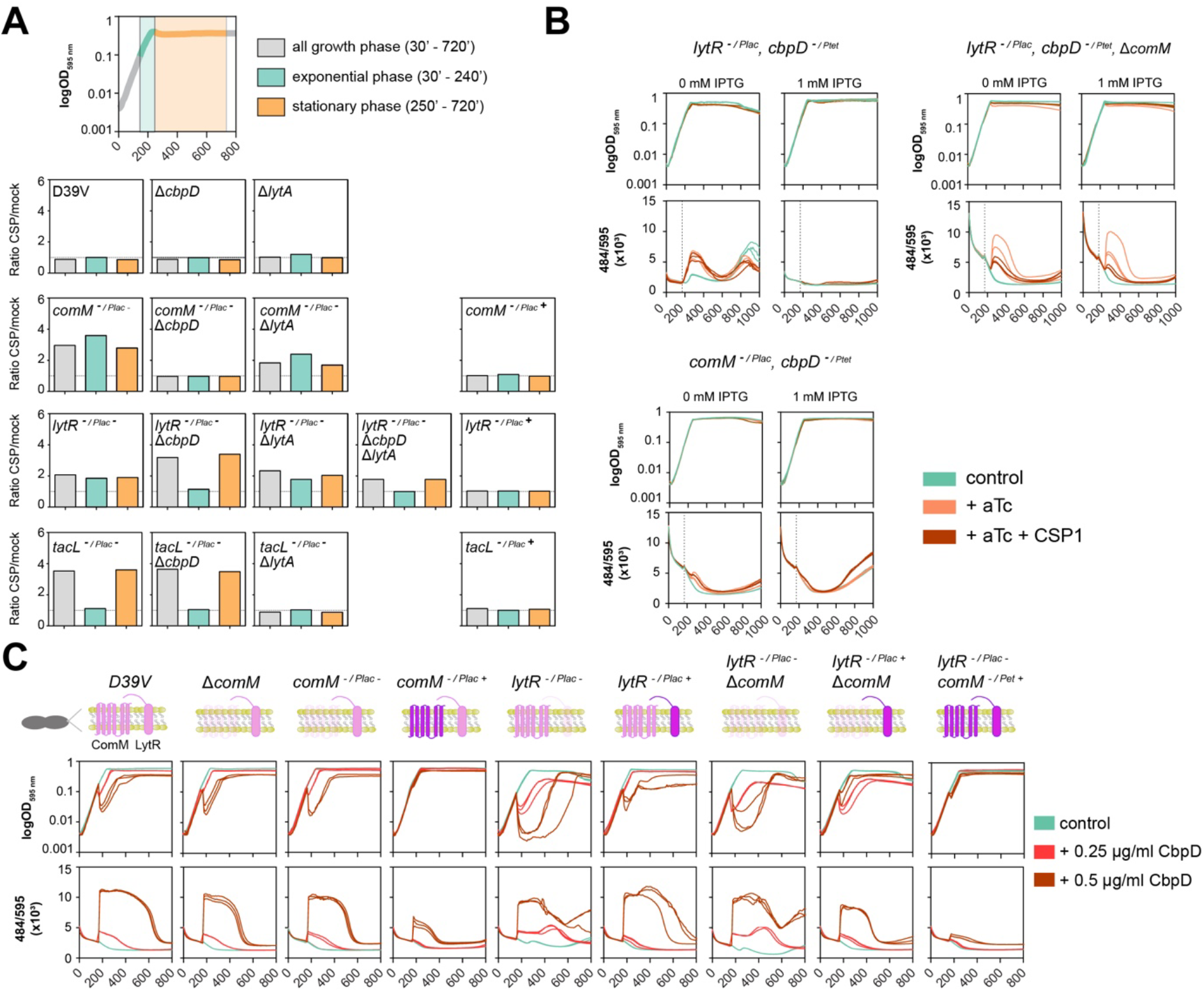
Both LytR and ComM are required for immunity to CbpD. **A)** Cell lysis evaluation in presence and absence of the fratricins. Top: growth phase was split in three periods: exponential phase (green; from CSP_1_ addition to stationary phase), stationary phase (orange) and all growth data across the curve (grey). Individual strains were grown in C+Y pH 6.8 to avoid natural competence development in presence of SYTOX™ Green Dead Cell Stain dye. When cell cultures reached OD_595 nm_ 0.1 (~ after 170 min), 0 and 100 ng/ml CSP_1_ was added to the medium, and the fluorescence ratio (+CSP_1_ / mock) was calculated. Three biological replicates per condition are shown. As fratricide occurs shortly after competence induction, it should be detected in the first period, while autolysis via LytA is detected during stationary phase. Strains containing a complementing copy are indicated by Plac (-indicates no IPTG, + indicates addition of 100 μM IPTG). **B)** Induced cell lysis by overexpression of *cbpD* (cbpD^-/Ptet+^). Individual strains were grown in C+Y pH 6.8 to avoid natural competence development in the presence of SYTOX™ Green Dead Cell Stain dye. When cell cultures reached OD_595 nm_ 0.1 (~ after 170 min), 0.5 μg/ml of anhydrotetracycline (aTc; orange) or 0.5 μg/ml of aTc plus 100 ng/ml of CSP_1_ (red) was added to induce *cbpD* expression and/or competence. Three biological replicates per condition are shown. **C)** Induced cell lysis by addition of recombinant CbpD (obtained as described in (Straume et al., 2020). Individual strains were grown in C+Y pH 6.8 to avoid natural competence development in presence of SYTOX™ Green Dead Cell Stain dye. When cell cultures reached OD_595 nm_ 0.1 (~ after 170 min), 0.25 μg/ml (red) or 0.5 μg/ml of purified CbpD was added to the medium. Three biological replicates per condition are shown.

### ComM is required for LytR activity

Competence activation, and subsequent upregulation of CbpD, ComM and LytR, might change the ratio between LTA and WTA (Figure 1A, 2A) as well as blocking septal PGN synthesis (Bergé et al., 2017) leading to protection from the PGN hydrolytic activity of CbpD. Indeed, cells become elongated when competence is induced and this depends on ComM (Figure S7, (Bergé et al., 2017; Straume et al., 2017a)). During exponential growth, the action of TacL ensures plentiful production of LTAs (Flores-Kim et al., 2019) (Figure 2A). To investigate this further, we measured the incorporation of TAs in the membrane and cell wall using radioactive Methyl-^3^H-Choline, as it is taken up and incorporated in TAs (Tomasz, 1967) (Figure 4A).

**Figure 4.**
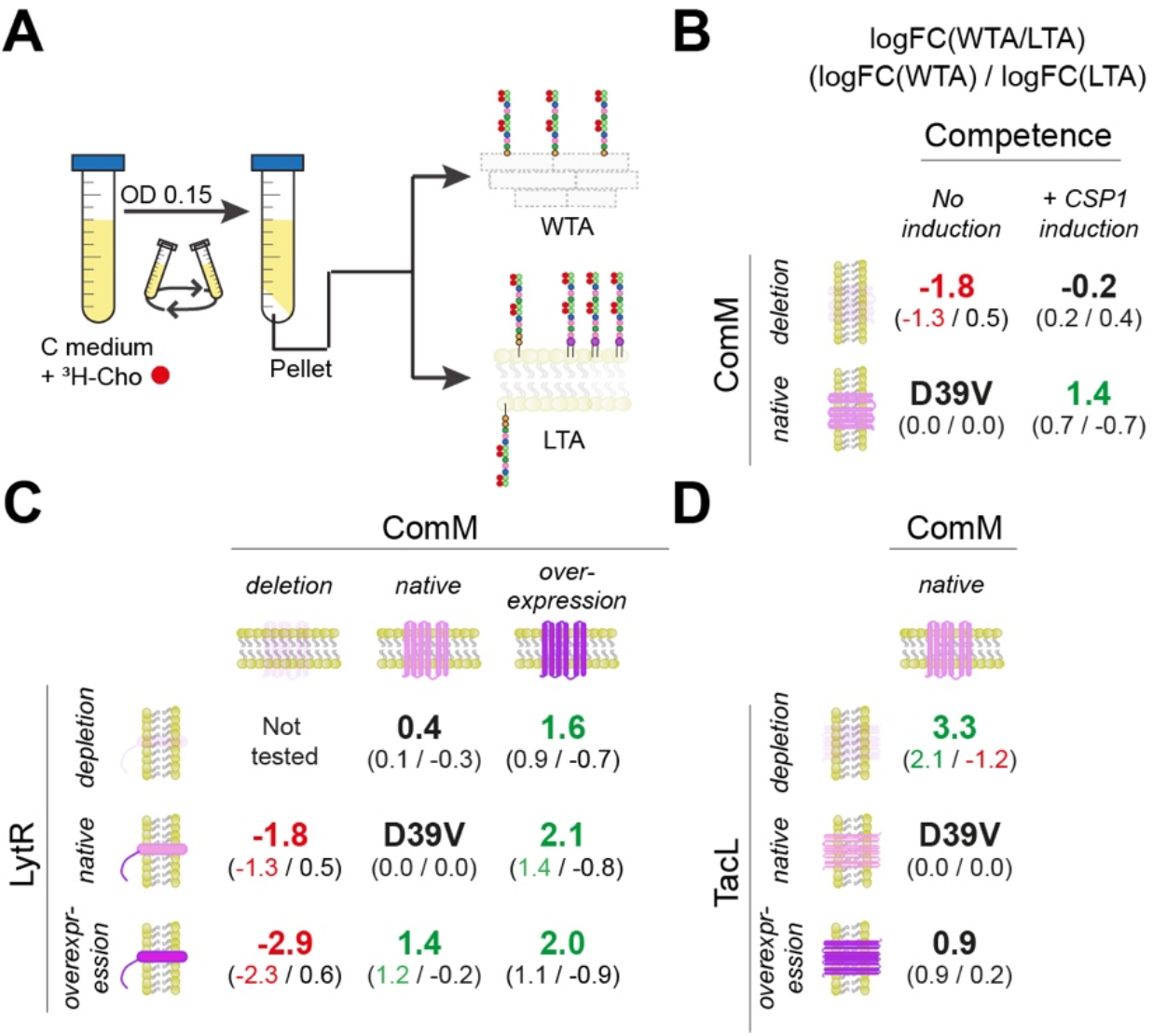
Competence induction leads to relatively more ^3^H-Cho incorporation in wall teichoic acids compared to lipoteichoic acids. **A)** Protocol to detect ^3^H-Cho. Cells were grown in C medium supplemented with ^3^H-Cho until OD_595 nm_ 0.15. The pellet was divided in two equally sized aliquots, one for membrane (including LTAs and TA precursors) and one for cell wall isolation (see methods section for more details). **B)** ^3^H-Cho detection after competence induction. Membranes and cell walls were isolated to quantify the amount of ^3^H-Cho incorporated. Average of three independent biological replicates and two technical replicates is shown (raw data in Table S3). In large the ratio in log fold change between WTA and LTA normalized to D39V wt cells (logFC(WTA0/logFC(LTA) is shown. In small is shown the relative abundancy of WTA or LTA compared to D39V (stated as 0.0/0.0). Red colour indicates a significantly smaller ratio between WTA/LTA in that condition compared to wt, whereas green colour indicates a significantly greater ratio between WTA/LTA. **C)** ^3^H-Cho detection in different LytR and ComM conditions. The condition with double *lytR* depletion and *comM* deletion was not tested due to the growth defect and increased lysis of the strain. LytR was depleted for ~ 2h to not cause too much cell lysis due to its essentiality. Average of three independent biological replicates and two technical replicates is shown (raw data in Table S2). **D)** ^3^H-Cho detection in different TacL conditions. Average of three independent biological replicates and two technical replicates is shown (raw data in Table S2).

Twenty minutes after competence induction in D39V, cell walls were separated from the membrane fraction (see Methods) and a significant increase in the WTA/LTA ratio was observed (Figure 4B, Table S3). This shift was abolished when competence was induced in absence of ComM (we used the Δ*comM* Δ*cbpD* double mutant to avoid fratricide, Figure S8A). In this strain, *comM* was replaced by a promoterless kanamycin-resistance cassette in frame, thus, levels of LytR are still highly increased after CPS1 addition. Indeed, an increase in WTA levels was observed in the *ΔcomM ΔcbpD* mutant strain when competence was triggered; however, those levels are significantly lower compared to D39V (Figure 4B) confirming that ComM is important for WTA production.

To analyse the role of both ComM and LytR in more detail, we tested ^3^H-Cho incorporation in strains with different expression levels of both genes (Figure 4C). Interestingly, *comM* overexpression resulted in a reduction of LTA amounts relative to WTA, independent of the *lytR* expression levels (Figure 4C). On the contrary, in absence of *comM* or in its native expression, the amounts of LTA remained similar to those in D39V. When cells were slightly depleted for LytR in native (low) *comM* levels, no significant differences were observed (Figure 5C). In addition, overexpression of LytR also resulted in increased amounts of WTA under basal ComM level conditions (Figure 4C, Table S3).

**Figure 5.**
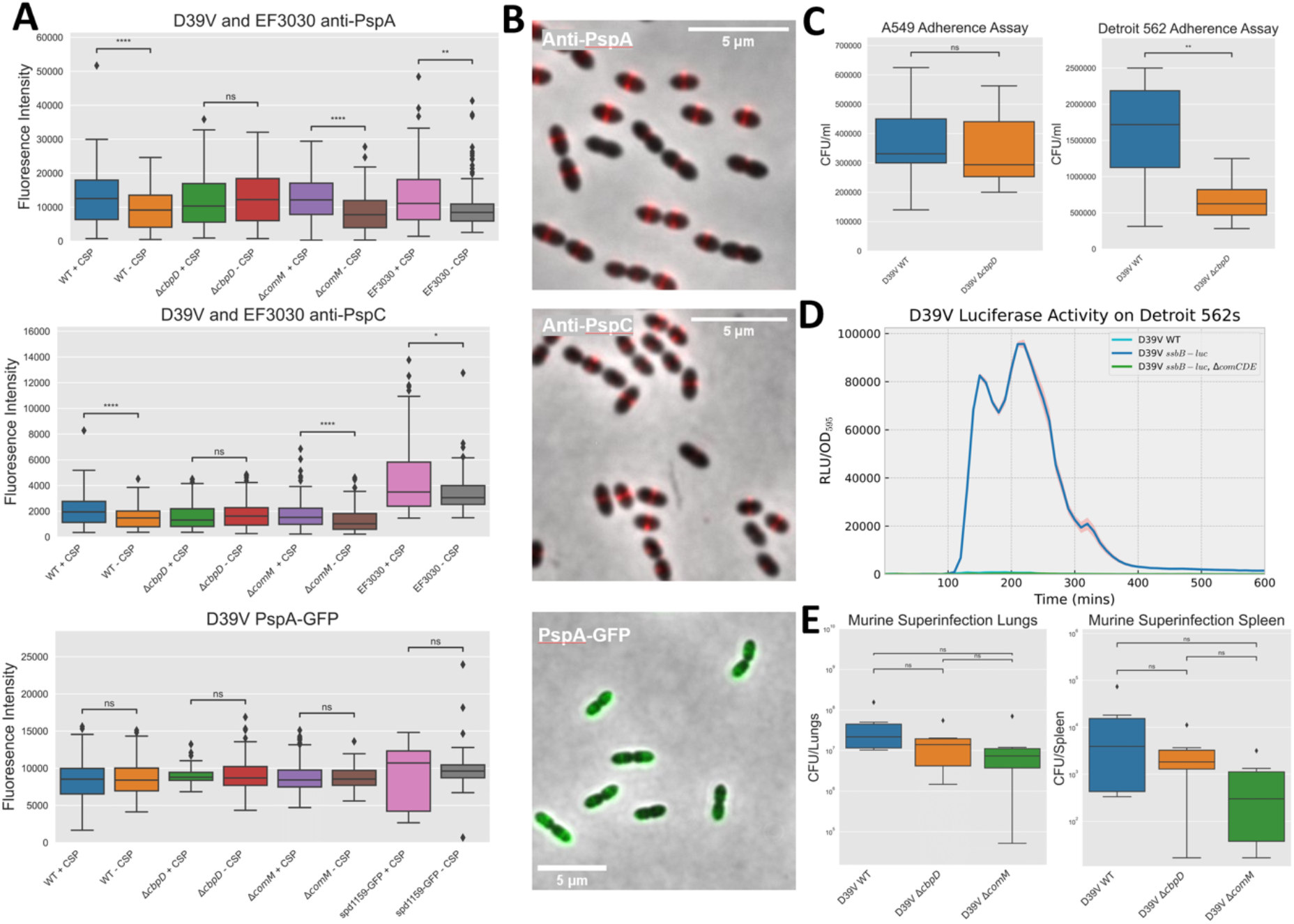
CbpD-dependent increased surface exposure of PspA and PspC, and impact during infection. **A)** D39V (WT, *ΔcbpD* and *ΔcomM*) and EF3030 cells, or corresponding D39V strains containing PspA genetically tagged with GFP to its C-terminus, were grown to OD 0.1 in C+Y medium pH 6.9, then exposed to 100 ng/ml or 0 ng/ml of CSP_1_ for 30 min. For the anti-PspA and anti-PspC microscopy, cells were then stained with primary antibodies raised against PspA or PspC, and then Goat anti-rabbit IgG (H+L) Alexa 555, both at 1/500 dilutions and subjected to epifluorescence microscopy. For the PspA-GFP microscopy, cells were subjected to epifluorescence microscopy directly after exposure to CSP_1_ for 30 mins. A GFP tagged SPV_1159 D39V strain was used a control as this gene’s regulation is not affected by competence induction (Kurushima et al., 2020). Fluorescence intensity based on phase-contrast and fluorescence composite images were measured. Diamond symbols represent outlier individual cells. Asterisks show statistically significant differences in fluorescence intensity (ns, not-significant, *, *P* < 0.05, **, *P* < 0.01, and ****, *P* < 0.0001, Mann-Whitney U test) (see methods section for more details). **B**) Representative phase contrast and fluorescence composite images taken of D39V WT after anti-PspA and anti-PspC immunostaining, and for D39V PspA-GFP after exposure to CSP_1_. **C**) Adherence assay. D39V (WT and *ΔcbpD*) strains were inoculated with A549 cells or Detroit 562 cells to a multiplicity of infection of ~20 in RPMI 1640 without phenol red supplemented with 1% (v/v) FBS and 10mM HEPES buffer for 2 h at 37°C. Cells were then detached, and appropriate dilutions of the cultures were plated on blood agar plates to determine the number of adherent bacteria (see methods section for more details). Data presented are the means ± standard deviation (ns, not-significant, **, *P* < 0.01, unpaired t-test). **D**) D39V (WT, *ssbB-luc, ssbB-luc* + Δ*comCDE*) cells were added to Detroit 562 nasopharyngeal cells to an OD_595_ 0.004 in C+Y pH 7.8 +/− 0.05 (permissive conditions for natural competence induction) containing 0.45 mg/ml of luciferine. Cells were incubated in 96-wells microtiter plates with no shaking. Growth (OD_595 nm_) and luciferase activity (RLU) were measured every 10 min for 14 h using a Tecan Infinite F200 PRO. An average of four replicates and the ± standard deviation are shown (see methods section for more details). **E**) Superinfection with D39V WT, *ΔcbpD* and *ΔcomM* mutants. Mice were infected intranasally with 50 plaque-forming units of the influenza A virus strain. Seven days later, mice were inoculated intranasally with 5×10^4^ CFU of *S. pneumoniae* strain D39V. Mice were sacrificed 24 hours post-infection and lungs and spleen were sampled to determine the bacterial viable counts. Data presented are the means ± standard deviation (ns, not-significant, unpaired t-test) from 6 replicates.

To complement these results, we also tested the depletion and overexpression of *tacL* (Figure 4D). In absence of TacL, TAs cannot be anchored in the membrane, thereby providing more substrate for LytR to produce WTA (Figure 2A). In support of this model, we find a shifted ratio of WTA/LTA in favour of WTA when TacL was depleted from the cells (Figure 4D). However, in the membrane fraction we still detected the TAs that are not yet anchored in the cell wall, in addition to the precursors anchored in the inner part of the membrane (Figure 4A). Overexpression of TacL did not impact LTA production during exponential growth, likely because the membrane is already saturated with LTAs in this growth phase (Flores-Kim et al., 2019) (Figure 4D, Table S3).

### Competence-dependent expression of CbpD exposes PspA and PspC to the cell surface

As shown above, upon competence activation, ComM and LytR alter the pneumococcal cell wall by changing the flux towards WTA biosynthesis and elongating the cell. Indeed, using phase-contrast microscopy, elongation of the cell upon competence induction was diminished in a Δ*comM* mutant (Figure S7A). In addition, the PGN hydrolase activity of CbpD would result in shedding of the capsule thereby liberating anchor sites on the PGN to which WTA could be attached by LytR. This would imply that choline-binding proteins, normally bound to the LTA, now have more binding sites available on the choline residues present on the WTA. To test whether CBPs indeed become more surface exposed during competence development, we performed immunostaining followed by fluorescence microscopy using antibodies raised against PspA and PspC. PspA and PspC are CBPs that play various roles during virulence from blocking complement deposition to binding to lactoferrin (Subramanian et al., 2019; Weiser et al., 2018). As shown in Figure 5A, activation of competence by the addition of synthetic CSP significantly increased antibody staining of both PspA and PspC as measured by immunofluorescence microscopy in strain D39V. Similar results were obtained with strain 19F, a recent clinical isolate (Junges et al., 2019). Crucially, this increased signal was abolished in a *cbpD* mutant but not in a *comM* mutant (Figure 5A). Complementary results were observed when over expressing CbpD or ComM. Here, only the increased expression of CbpD led to higher antibody staining of PspA (Figure S7B). Surprisingly, immunofluorescence microscopy showed a clear septal localization for both PspA and PspC (Figure 5B), while ComM genetically tagged with a yellow fluorescent protein also demonstrated an enriched localization pattern at the septum (Figure S7C). This is in line with previous findings, where LytR was previously also shown to be enriched at the septa and CbpD was shown to bind there as well (Eberhardt et al., 2012; Eldholm et al., 2010). Additionally, localisation of PspA fused to GFP showed a more uniform cell wall/membrane localization, whose signal did not change during competence (Figure 5A). In corroboration, immunofluorescence microscopy using an encapsulated mutant displayed less septal localisation and more background fluorescence (Figure S7D). These results suggest that while PspA is present throughout the pneumococcal cell surface, it is mainly surface exposed at the newly synthesized septum (Figure 6). Moreover, these data indicate that CBPs become more surface exposed during competence development and that this depends on the activity of CbpD. In line with these results, a *cbpD* mutant has been shown to be significantly attenuated in its ability to colonise the nasopharynx of rats (Gosink et al., 2000). Moreover, adherence assays of D39V WT and corresponding *cbpD* deletion mutant indicated that absence of CbpD attenuated pneumococcal adherence to human nasopharyngeal Detroit 562 cells, but not for A549 lung epithelial cells (Figure 5C). A luciferase assay using a competence-specific reporter demonstrated that competence is indeed activated in the Detroit 562 adherence assay (Figure 5D). Moreover, in a murine influenza A virus superinfection model, we showed that both *cbpD* and *comM* deletion mutants were not significantly attenuated during infection in the lungs or spleen (Figure 5E). This suggests that CbpD is important for colonization but not strictly during replication in the context of flu. These data support a model in which competence is important for pneumococcal adherence by shuttling key virulence proteins, the CBPs, to the outside surface of the cell to interact with the host (Figure 6).

**Figure 6.**
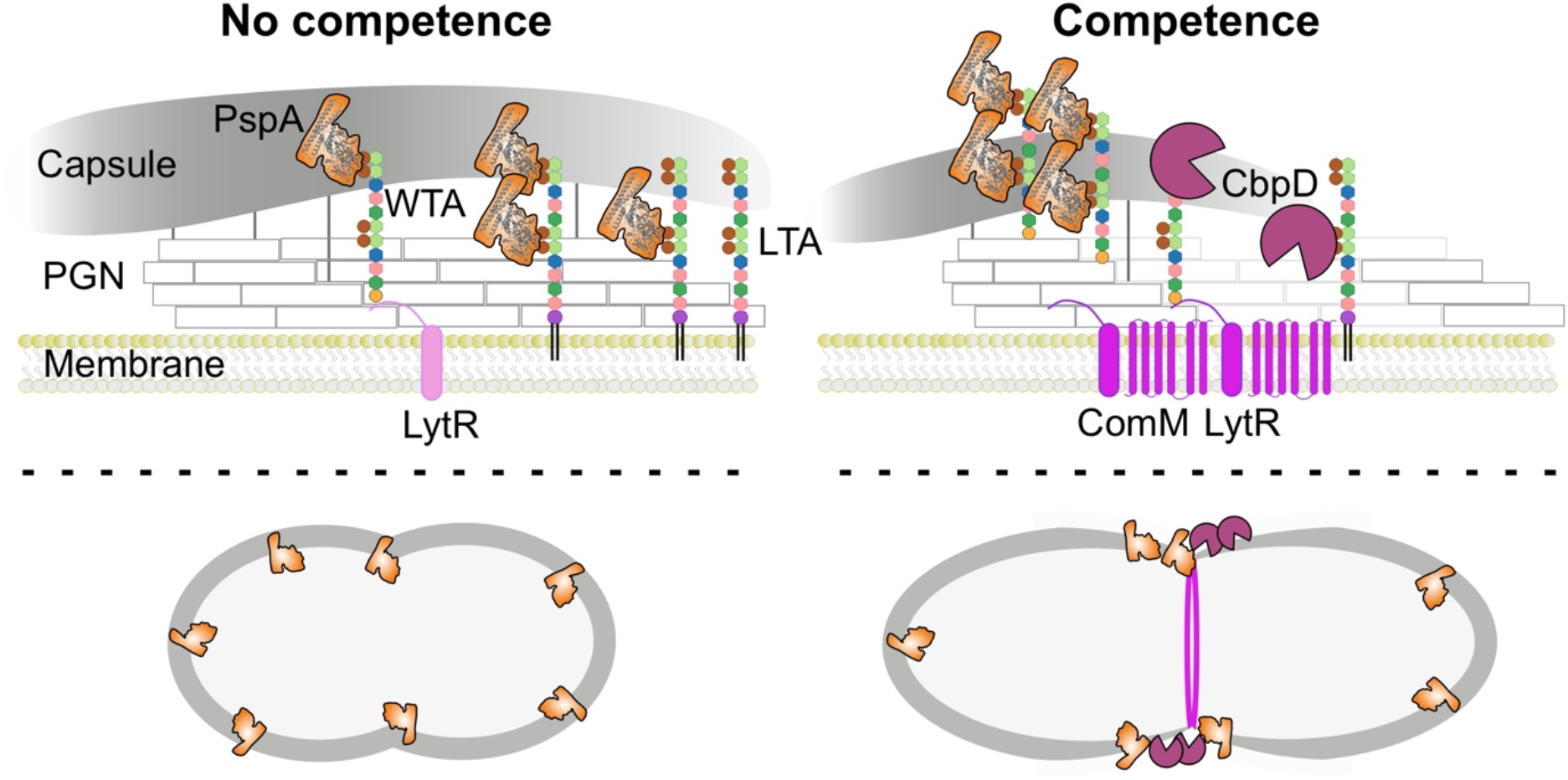
Model for competence-dependent surface exposure of key virulence factors. During normal growth, basal levels of LytR support growth by the anchoring of teichoic acid precursor to peptidoglycan (PGN) to produce wall teichoic acids (WTA) (left situation). Cells are fully covered by the capsule under non-competent conditions. Note that the choline-containing WTA is competing with the capsule for binding to PGN at the same sites on the PGN. Subsequently, choline-binding proteins, including virulence factor PspA, are poorly surface exposed. During competence (right situation), ComM and CbpD are produced, as well as more LytR that is located in the same operon as *comM* (Figure 2). Activation of competence results in cell elongation through ComM localized at the septum (Bergé et al., 2017). Together, this results in a shift in presence of WTA vs lipoteichoic acid (LTA), providing relatively more choline residues available for binding by PspA within the cell wall (Figure 5). The process is augmented by cleavage of peptidoglycan (PGN) at midcell by CbpD, but cell lysis is prevented through ComM. Together, this results in thinning of capsule at midcell, further exposing PspA to the bacterial cell surface. This model could explain why CbpD mutants show reduced colonization and why competence mutants in general are less virulent (see Discussion).

## Discussion

In our study we identified genes that become essential during competence in the human pathogen *S. pneumoniae*. Using CRISPRi-seq we found 14 sgRNAs (targeting 10 operons) underrepresented during competence. Two competence-related operons, *comCDE* and *comM*, showed an expected fitness cost as the immunity protein ComM cannot be produced by downregulation of both operons, and thus fratricins produced by competing bacteria in the pool lyse these cells. Contrary, the *comAB* operon did not show any fitness cost as cells can still sense exogenous CSP produced by competing cells and activate competence (Figure S2B). The fact that the sgRNA targeting *comX* was present in all conditions, confirms that fratricide immunity is conferred due to ComM (which is ComE-P dependent and not ComX dependent), rather than other competence-related proteins (Figure S2B).

Interestingly, most of the targeted operons were related to the TA synthesis pathways, suggesting their important role protecting cells from competence-related autolysis (Figure 1B). Other genes known to play a role in cell wall synthesis were also underrepresented during competence development but did not make the statistical cut-off (Table S1). This suggests that perturbations in general cell wall homeostasis also sensitizes pneumococci to fratricide. Indeed, it was previously shown that cells depleted for *pbp2b, rodA, mreD, divIVA* and *cozEa* are more susceptible to lysis by CbpD (Straume et al., 2017b). Contrary to other Gram-positive bacteria, the TAs biosynthesis pathway in *S. pneumoniae* is identical for LTAs and WTAs (Denapaite et al., 2012; Seo et al., 2008). Using radiolabeled choline, we show that the levels or relative amounts of LTA and WTA are altered when competence is triggered.

At the heart of immunity towards lysis by CbpD lies ComM (Figure 1A, 1C). Here, we show that ComM works in concert with LytR, and expression of both genes provides optimal immunity towards CbpD (Figure 3C). This work also provides strong evidence that LytR is the key enzyme of the LCP-protein family member that anchors the TA precursor to PGN to create WTA (Figure 4). Based on these data, we propose a model in which the flux of TA from LTA increases towards WTA during competence development by the upregulation of *comM* and *lytR* (Figure 6). This will aid *de novo* formed CBPs to bind to nascent WTA at the midcell. Alternatively, or at the same time, the action of CbpD will result in shedding of the capsule thereby liberating WTA binding sites on the PGN. Together, the competence-dependent remodelling of the cell wall leads to increased surface exposure of CBPs that include major virulence factors such as PspA and PspC (Figure 6). Using immunofluorescence, we show that both PspA and PspC are surface exposed mainly at the midcell, both in serotype 2 strain D39V and in a serotype 19F strain (Figure 5). In this respect, it is interesting to note that CbpD, ComM and LytR also show septal localization (Figure 5C and (Eberhardt et al., 2012), suggesting that the cell wall is predominantly remodelled at the septum during competence and that these provide the highest affinity sites for CBPs. Whether these altered TA fluxes are indirect effects of general ComM-mediated cell wall remodelling, through for instance StkP (Bergé et al., 2017), or that the major function of ComM is to redirect TA to WTA and that this causes upstream effects on septal PGN synthesis remains to be determined. Interestingly, it was recently shown that CbpD only attacks PGN that is newly synthesized by PBP2x and FtsW (Straume et al., 2020). Thus, increased WTA-levels (mediated by ComM and LytR during competence) could somehow inhibit the activity of PBP2x or other essential components of the divisome, thereby making the cells resistant to CbpD. This idea is also in line with the observation that non-competent pneumococci depleted for TacL, and thus with increased levels of WTA (Fig 4D; (Flores-Kim et al., 2019), are resistant to fratricide. The subsequent delay in division might be important to resolve chromosome dimers that can occur during transformation (Bergé et al., 2017).

Why would competence for genetic transformation be required during virulence? Having access to new DNA is not of direct benefit during acute infection. It is tempting to speculate that cell surface remodelling via CbpD-ComM-LytR is one of the main reasons that competence development is switched on during infection. If even a small pool of the CBPs, many of which are bona fide virulence factors such as CbpN (aka PcpA), CbpC, CbpG, PspA and PspC (Gosink et al., 2000; Hakenbeck et al., 2009; Subramanian et al., 2019; Weiser et al., 2018) become more surface exposed, then this might allow bacteria to better adhere or evade the host’s immune system (Figure 6). In addition, an increased flux in WTA would also imply reduced capsule anchoring, further improving adherence and surface exposure of the CBPs (Selinger and Reed, 1979; Weiser et al., 2018). Whether competence development indeed affects capsule shedding at midcell remains to be tested. Altogether, the data presented here could explain why CbpD mutants were observed to be attenuated in adherence to human nasopharyngeal cells (Fig. 5C) and colonization of the rat nasopharynx (Gosink et al., 2000). Interestingly, a *cbpD* mutant was not altered in its ability to adhere to A549 lung epithelial cells, likely because adherence is already poor on these cells with wild type bacteria (Fig. 5C). Interestingly, while a *cbpD* mutant was not attenuated in its ability to replicate in a mouse viral superinfection model (Fig. 5E), *comCDE* mutants were shown to be attenuated in a mouse and zebrafish meningitis model. The key difference between these models is that in the superinfection model bacteria can immediately replicate without a clear bottleneck imposed by the host immune system (Liu et al., 2021), while the meningitis models take significant time from inoculation to colonization to dissemination (Jim et al., 2022; Schmidt et al., 2019). It will be interesting to see how competence and ComM-LytR regulated WTA synthesis plays a role in other disease models as well as in other clinical strains.

## Acknowledgements

We thank Olivier Bützberger for technical assistance, construction of strains, and CRISPRi-seq data collection. We also thank Thomas Kohler (University of Greifswald) for supervision and generating antibodies. Work in the Veening lab is supported by the Swiss National Science Foundation (SNSF) (project grants 310030_200792 and 310030_192517), SNSF JPIAMR grant (40AR40_185533), SNSF NCCR ‘AntiResist’ (51NF40_180541) and ERC consolidator grant 771534-PneumoCaTChER. This work was further supported by grants of the Deutsche Forschungsgemeinschaft to N.G. (GI 979/1-2), S.H. (HA 3125/5-2). Work in the Perez lab is supported by the SNSF (PP00P3_198903) and the Helmut Horten Stiftung (HHS).

## Methods

### STAR methods

#### LEAD CONTACT AND MATERIALS AVAILABILITY

Further information and requests for resources and reagents should be directed to and will be fulfilled by the Lead Contact, Jan-Willem Veening.

This study did not generate new unique reagents.

#### EXPERIMENTAL MODEL AND SUBJECT DETAILS

##### Bacterial strains and growth conditions

All pneumococcal strains used in this study are derivatives of the clinical isolate *S. pneumoniae* D39V (Slager et al., 2018), Public Health England NCTC14078). Bacterial strains are listed in Table S2. Growth conditions of bacterial cells were described previously (Domenech et al., 2020). Briefly, *S. pneumoniae* was grown in C+Y medium (pH 6.8, non-permissive conditions for natural competence induction), at 37 °C and stored at –80 °C in C+Y with 14.5% glycerol at OD_595 nm_ of 0.4.

#### METHOD DETAILS

##### Competence assays

Competence development was monitored in strains containing a transcriptional fusion of the firefly luciferase gene (*luc*) with the late competence gene *ssbB*. The *S. pneumoniae* strains were cultured in a Tecan Infinite F200 PRO allowing for real-time monitoring of competence induction *in vitro*. A pre-culture was diluted to initial OD_595nm_ 0.004 in C+Y pH 7.8 +/− 0.05 (permissive conditions for natural competence induction) containing 0.45 mg/ml of luciferine and then incubated in 96-wells microtiter plates with no shaking. Growth (OD_595 nm_) and luciferase activity (RLU) were measured every 10 min for 14 h. Expression of the *luc* gene (only if competence is activated) results in the production of luciferase and thereby in the emission of light (Domenech et al., 2020). An average of three replicates and the standard error of the mean (SEM) are shown unless indicated.

##### Pooled CRISPRi library

The pooled CRISPRi library containing 1499 sgRNAs targeting all the transcription start sites in *S. pneumoniae* D39V strain (Liu et al., 2021) was used to identify essential genes during competence. A pre-culture of the pooled library was grown at in C+Y medium pH 6.8 to avoid natural competence induction, at 37 °C until OD_595 nm_ of 0.1 (Figure 1A). Then, the pre-culture was diluted at OD_595 nm_ 0.005 in presence or absence of 1 mM IPTG to induce the expression of *dCas9*. After 2 h (OD_595 nm_ ~ 0.1) of library induction, 100 ng/ml CSP_1_ was added in half of the samples to induce competence. Cells were incubated at 37 °C until OD_595 nm_ 0.4, and DNA was isolated using a commercial kit (Promega®). Preparation of the Ilumina library by one-step PCR with commercial oligos and library sequencing were performed following the manufacturer instructions and described previously (de Bakker et al., 2022).

##### Construction of inducible strains

For every gene related to teichoic acid synthesis, an ectopic IPTG-inducible strain was created using the pASR130 (pPEPZ::P*_lac_*-TmpR-blaR) plasmid that integrates at the non-essential ZIP locus (Keller et al., 2019). Primers used to amplify the genes are listed in Table S4. PCR products were digested with BsmBI, BsaI or SapI (depending on the presence of restriction sites incompatible with the cloning) and were ligated with similarly digested pPEPZ plasmid containing the P*_lac_* promoter. The ligation was transformed into strain ADP95 (D39V, *prs1::PF6-lacI-tetR, bgaA::P_ssbB_-luc*; (Domenech et al., 2020). All transformants were selected on Columbia blood agar containing 10 μg/ml of trimethoprim and correct colonies were verified by PCR and sequencing.

##### Construction of the native deletions

Every gene related to teichoic acid synthesis was replaced by an erythromycin-resistant marker, in frame with the rest of the genes in the same operon (without promoter and terminator). Primers used to amplify the genes are listed in Table S4. Upstream region (~ 1Kb), downstream region (~ 1Kb) and the promoterless erythromycin marker were digested with the indicated restriction enzyme (BsmBI or BsaI, depending on the presence of restriction sites incompatible with the cloning) and were ligated. The ligation was transformed into the corresponding strain carrying the ectopic inducible construct. All transformants were selected on Columbia blood agar containing 0.5 μg/ml of erythromycin and 1 mM IPTG, and correct colonies were verified by PCR and sequencing.

##### In vivo *radiolabelling of TA*

*S. pneumoniae* strains (D39V derivatives) were grown in C medium at pH 6.8 (to avoid natural competence development) until OD_595 nm_ of 0.1 in absence of the inducers. The pre-cultures were diluted to initial OD_595 nm_ of 0.001 in 10 ml of C medium at pH 6.8 and supplemented with 43.7 mM of ^3^H-choline. Final concentrations of 1 mM of IPTG and/or 500 ng/ml of aTc were added when indicated. Cells were grown at 37 °C without shaking until OD_595 nm_ of 0.15. For the conditions that required competence induction, 100 ng/ml of CSP_1_ was added at OD_595 nm_ 0.10 and incubated for 20 minutes at 37 °C. Cells were centrifuged for 5 min at 7000 x *g* and 2 ml of supernatant was collected and stored at −80°C. The pellet was resuspended in 2 ml of 50 mM MES and distributed in two fractions. Membrane and cell wall isolates were obtained as described before (Flores-Kim et al., 2019). Each growth condition was performed with experimental triplicates.

##### Radioactivity quantification

Radioactivity of cell wall and membrane isolates was measured by scintillation counting using a Liquid Scintillation Analyzer Tri-Carb 4910 TR (PerkinElmer). To do this, 2.5 ml of Ultima Gold™ XR LSC Cocktail (PerkinElmer, Waltham, MA) were mixed either with 100 ml of isolated membrane or 200 ml of cell wall isolate. Each experimental triplicate was measured twice. For background correction, 500 ml of cell culture supernatant was treated and measured in the same way.

###### Immunostaining followed by microscopy

*S. pneumoniae* cells were grown in C+Y medium pH 7.4 at 37 °C to an OD_595_ = 0.1 or in RPMI1640. For microscopy: Upon reaching OD_595_ = 0.1, 100ng/ml of CSP-1 and/or 0.5μg/ml aTc was added and cells were left for 30 min at 37°C before centrifugation at 10,000 g for 3 min. Cells were then washed with 1 ml of 1x PBS before incubation with PspA or PspC antibodies at a 1/500 dilution in 100μl 1x PBS at 37°C for 1h. Cells were again centrifuged and washed with 1ml of 1x PBS before incubation with 1/500 diluted Goat anti-rabbit IgG (H+L) Alexa 555 (Thermo Fisher Scientific) in 100μl 1x PBS at 37°C for 1h. Cells were then washed twice with 1ml of 1x PBS and resuspended in 50 μl of 1x PBS. 0.4 μl of cells were subsequently spotted onto PBS-polyacrylamide (10%) pads within a gene-frame (Thermo Fisher Scientific) and sealed with a cover slip as described previously (de Jong et al., 2011). Microscopy acquisition was performed using a Leica DMi8 microscope with a sCMOS DFC9000 (Leica) camera and a SpectraX light source (Lumencor). Phase-contrast images were acquired using transmission light (100 ms exposure) and still fluorescence images were acquired with 700 ms exposure. The Leica DMi8 filters set used were as followed: YFP (Ex: 500, Dc: 520, Em: 535), Alexa 555 (Ex: 550, Dc: 570, Em: 576) and GFP (Ex: 470/40 nm Chroma ET470/40x, BS: LP 498 Leica 11536022, Em: 520/40 nm Chroma ET520/40 m). Images were processed using LasX v.3.4.2.18368 (Leica). All microscopy images were processed using FIJI v.1.52q (fiji.sc). Fluorescence intensity based on phase-contrast and fluorescence composite images, as well as cell length measurements, were performed using MicrobeJ (Ducret et al., 2016).

##### Detroit 562 adherence and competence assays

Detroit 562 nasopharyngeal and A549 lung epithelial cells were grown in Dulbecco’s modified Eagle’s medium (DMEM) supplemented with 10% foetal calf serum (FCS), 25mM HEPES buffer (Sigma-Aldrich) and 5ml of Penicillin-Streptomycin (5,000 U/mL) (Gibco™), in 75 cm2 tissue culture flasks (Corning®) at 37°C in a 5% CO2 atmosphere. 1ml of 4×10^5^ of Detroit 562 or 2×10^5^ A549 cells in the DMEM media were seeded into each well of a 24-well tissue culture tray (Corning Costar) and incubated at 37°C, 5% CO2 for 24h. Cells were then washed with 1ml 1x PBS and 1ml of RPMI 1640 medium without phenol red (Gibco™) supplemented with 10% FCS for each well was added. The following day, pneumococcal strains VL1 and VL561 were grown to an OD of 0.1 in C+Y media before adding 0ng/ml or 100ng/ml and incubating for 30 min. Bacteria were then spun down and resuspended in infection medium, RPMI 1640 without phenol red supplemented with 1% (v/v) FBS and 10mM HEPES buffer. Bacterial samples were added to wells in the 24 well tray containing cells to a multiplicity of infection of ~20, and incubated at 37°C, 5% CO2. After 2 h, samples were washed twice with 1ml 1xPBS, and cells were detached from the plate by treatment with 100μl of 15mM sodium citrate and 400μ 0.1% Triton x-100. Appropriate dilutions of the cultures were plated on blood agar plates to determine the number of adherent bacteria. Assays were performed in quadruplicate from two independent experiments. For the pneumococcal D39V competence assay on nasopharyngeal cells, each well of a 96 well tray (Corning costar) was seeded with 200μl of 5×10^5^ of Detroit 562 cells in the DMEM media and incubated at 37°C, 5% CO2 for 24h. Cells were then washed with 200μl 1x PBS and 200μl of RPMI 1640 medium without phenol red (Gibco™) supplemented with 10% FCS for each well was added. The following day, the competence assay was performed as described above on D39V WT and corresponding strains containing a *ssbB*-*luc* and/or *cbpD* deletion mutations. An average of four replicates and the standard deviation (SD) are shown.

##### Murine Superinfection

Male C57BL/6JRj mice (8 weeks old) (Janvier Laboratories, Saint Berthevin, France) were maintained in individually ventilated cages and were handled in a vertical laminar flow cabinet (class II A2, ESCO, Hatboro, PA). All experiments complied with national, institutional and European regulations and ethical guidelines, were approved by our Institutional Animal Care and Use guidelines (D59-350009, Institut Pasteur de Lille; Protocol APAFIS#16966 201805311410769_v3) and were conducted by qualified, accredited personnel. Mice were anesthetized by intraperitoneal injection of 1.25 mg (50 mg/kg) ketamine plus 0.25 mg (10 mg/kg) xylazine in 200 μl of PBS. Mice were infected intranasally with 30 μl of PBS containing 50 plaque-forming units (PFUs) of the pathogenic murine-adapted H3N2 influenza A virus strain Scotland/20/74 (Matarazzo et al., 2019). Seven days later, mice were inoculated intranasally with 5.10^4^ CFU of *S. pneumoniae* strain in 30 μl of PBS. Mice were sacrificed 24 hours post-infection by intraperitoneal injection of 5.47 mg of sodium pentobarbital in 100 μl PBS (Euthasol, Virbac, France). Lungs and spleen were sampled to determine the bacterial load. Tissues were homogenized with an UltraTurrax homogenizer (IKA-Werke, Staufen, Germany) and serial dilutions were plated on blood agar plates and incubated at 37°C. Viable counts were determined 24h later.

#### QUANTIFICATION AND STATISTICAL ANALYSIS

Data analysis was performed using GraphPad Prism, Microsoft Excel, R version 2.15.1 and RStudio Version 1.0.136.

Data shown in plots are represented as mean of at least three replicates ± SEM, as stated in the figure legends. Exact number of replicates for each experiment are enclosed in their respective figure legends.

## DATA AND CODE AVAILABILITY

The fastq files generated from sequencing are available on NCBI with accession number PRJNA841864. The 20 bp base-pairing sequences were trimmed, mapped and counted with 2FAST2Q (Bravo et al., 2021). The count data of sgRNAs were then analyzed with the DESeq2 package in R (https://github.com/veeninglab/CRISPRi-seq) for evaluation of fitness cost of each sgRNA. We tested against a log2FC of 1, with an alpha of 0.05.

## KEY RESOURCES TABLE

Supplementary File named Key_Resources_Table will be made available at a later stage.

## References

Aggarwal, S.D., Eutsey, R., West-Roberts, J., Domenech, A., Xu, W., Abdullah, I.T., Mitchell, A.P., Veening, J.-W., Yesilkaya, H., and Hiller, N.L. (2018). Function of BriC peptide in the pneumococcal competence and virulence portfolio. PLoS Pathog. 14, e1007328. https://doi.org/10.1371/journal.ppat.1007328.

Aprianto, R., Slager, J., Holsappel, S., and Veening, J.-W. (2016). Time-resolved dual RNA-seq reveals extensive rewiring of lung epithelial and pneumococcal transcriptomes during early infection. Genome Biol. 17, 198. https://doi.org/10.1186/s13059-016-1054-5.

Aprianto, R., Slager, J., Holsappel, S., and Veening, J.-W. (2018). High-resolution analysis of the pneumococcal transcriptome under a wide range of infection-relevant conditions. Nucleic Acids Res. 46, 9990–10006. https://doi.org/10.1093/nar/gky750.

de Bakker, V., Liu, X., Bravo, A.M., and Veening, J.-W. (2022). CRISPRi-seq for genome-wide fitness quantification in bacteria. Nat. Protoc. 17, 252–281. https://doi.org/10.1038/s41596-021-00639-6.

Bergé, M.J., Mercy, C., Mortier-Barrière, I., VanNieuwenhze, M.S., Brun, Y.V., Grangeasse, C., Polard, P., and Campo, N. (2017). A programmed cell division delay preserves genome integrity during natural genetic transformation in Streptococcus pneumoniae. Nat. Commun. 8, 1621. https://doi.org/10.1038/s41467-017-01716-9.

Bonnet, J., Durmort, C., Mortier-Barrière, I., Campo, N., Jacq, M., Moriscot, C., Straume, D., Berg, K.H., Håvarstein, L., Wong, Y.-S., et al. (2018). Nascent teichoic acids insertion into the cell wall directs the localization and activity of the major pneumococcal autolysin LytA. Cell Surf. Amst. Neth. 2, 24–37. https://doi.org/10.1016/j.tcsw.2018.05.001.

Bravo, A.M., Typas, A., and Veening, J.-W. (2021). 2FAST2Q: A general-purpose sequence search and counting program for FASTQ files. 2021.12.17.473121. https://doi.org/10.1101/2021.12.17.473121.

Chewapreecha, C., Harris, S.R., Croucher, N.J., Turner, C., Marttinen, P., Cheng, L., Pessia, A., Aanensen, D.M., Mather, A.E., Page, A.J., et al. (2014). Dense genomic sampling identifies highways of pneumococcal recombination. Nat. Genet. 46, 305–309. https://doi.org/10.1038/ng.2895.

Claverys, J.-P., Martin, B., and Polard, P. (2009). The genetic transformation machinery: composition, localization, and mechanism. FEMS Microbiol. Rev. 33, 643–656. https://doi.org/10.1111/j.1574-6976.2009.00164.x.

Croucher, N.J., Harris, S.R., Fraser, C., Quail, M.A., Burton, J., van der Linden, M., McGee, L., von Gottberg, A., Song, J.H., Ko, K.S., et al. (2011). Rapid pneumococcal evolution in response to clinical interventions. Science 331, 430–434. https://doi.org/10.1126/science.1198545.

Dagkessamanskaia, A., Moscoso, M., Hénard, V., Guiral, S., Overweg, K., Reuter, M., Martin, B., Wells, J., and Claverys, J.-P. (2004). Interconnection of competence, stress and CiaR regulons in Streptococcus pneumoniae: competence triggers stationary phase autolysis of ciaR mutant cells. Mol. Microbiol. 51, 1071–1086. https://doi.org/10.1111/j.1365-2958.2003.03892.x.

Dawson, M.H., and Sia, R.H. (1931). IN VITRO TRANSFORMATION OF PNEUMOCOCCAL TYPES: I. A TECHNIQUE FOR INDUCING TRANSFORMATION OF PNEUMOCOCCAL TYPES IN VITRO. J. Exp. Med. 54, 681–699. https://doi.org/10.1084/jem.54.5.681.

Denapaite, D., Brückner, R., Hakenbeck, R., and Vollmer, W. (2012). Biosynthesis of teichoic acids in Streptococcus pneumoniae and closely related species: lessons from genomes. Microb. Drug Resist. Larchmt. N 18, 344–358. https://doi.org/10.1089/mdr.2012.0026.

Domenech, A., Brochado, A.R., Sender, V., Hentrich, K., Henriques-Normark, B., Typas, A., and Veening, J.-W. (2020). Proton Motive Force Disruptors Block Bacterial Competence and Horizontal Gene Transfer. Cell Host Microbe 27, 544–555.e3. https://doi.org/10.1016/j.chom.2020.02.002.

Ducret, A., Quardokus, E.M., and Brun, Y.V. (2016). MicrobeJ, a tool for high throughput bacterial cell detection and quantitative analysis. Nat. Microbiol. 1, 16077. https://doi.org/10.1038/nmicrobiol.2016.77.

Eberhardt, A., Hoyland, C.N., Vollmer, D., Bisle, S., Cleverley, R.M., Johnsborg, O., Håvarstein, L.S., Lewis, R.J., and Vollmer, W. (2012). Attachment of capsular polysaccharide to the cell wall in Streptococcus pneumoniae. Microb. Drug Resist. Larchmt. N 18, 240–255. https://doi.org/10.1089/mdr.2011.0232.

Eldholm, V., Johnsborg, O., Haugen, K., Ohnstad, H.S., and Håvarstein, L.S. (2009). Fratricide in Streptococcus pneumoniae: contributions and role of the cell wall hydrolases CbpD, LytA and LytC. Microbiol. Read. Engl. 155, 2223–2234. https://doi.org/10.1099/mic.0.026328-0.

Eldholm, V., Johnsborg, O., Straume, D., Ohnstad, H.S., Berg, K.H., Hermoso, J.A., and Håvarstein, L.S. (2010). Pneumococcal CbpD is a murein hydrolase that requires a dual cell envelope binding specificity to kill target cells during fratricide. Mol. Microbiol. 76, 905–917. https://doi.org/10.1111/j.1365-2958.2010.07143.x.

Elm, C., Braathen, R., Bergmann, S., Frank, R., Vaerman, J.-P., Kaetzel, C.S., Chhatwal, G.S., Johansen, F.-E., and Hammerschmidt, S. (2004). Ectodomains 3 and 4 of human polymeric Immunoglobulin receptor (hpIgR) mediate invasion of Streptococcus pneumoniae into the epithelium. J. Biol. Chem. 279, 6296–6304. https://doi.org/10.1074/jbc.M310528200.

Flores-Kim, J., Dobihal, G.S., Fenton, A., Rudner, D.Z., and Bernhardt, T.G. (2019). A switch in surface polymer biogenesis triggers growth-phase-dependent and antibiotic-induced bacteriolysis. ELife 8, e44912. https://doi.org/10.7554/eLife.44912.

Galán-Bartual, S., Pérez-Dorado, I., García, P., and Hermoso, J.A. (2015). Chapter 11 - Structure and Function of Choline-Binding Proteins. In Streptococcus Pneumoniae, J. Brown, S. Hammerschmidt, and C. Orihuela, eds. (Amsterdam: Academic Press), pp. 207–230.

Gosink, K.K., Mann, E.R., Guglielmo, C., Tuomanen, E.I., and Masure, H.R. (2000). Role of novel choline binding proteins in virulence of Streptococcus pneumoniae. Infect. Immun. 68, 5690–5695. https://doi.org/10.1128/IAI.68.10.5690-5695.2000.

Hakenbeck, R., Madhour, A., Denapaite, D., and Brückner, R. (2009). Versatility of choline metabolism and choline-binding proteins in Streptococcus pneumoniae and commensal streptococci. FEMS Microbiol. Rev. 33, 572–586. https://doi.org/10.1111/j.1574-6976.2009.00172.x.

Håvarstein, L.S., Coomaraswamy, G., and Morrison, D.A. (1995). An unmodified heptadecapeptide pheromone induces competence for genetic transformation in Streptococcus pneumoniae. Proc. Natl. Acad. Sci. U. S. A. 92, 11140–11144. https://doi.org/10.1073/pnas.92.24.11140.

Håvarstein, L.S., Gaustad, P., Nes, I.F., and Morrison, D.A. (1996). Identification of the streptococcal competence-pheromone receptor. Mol. Microbiol. 21, 863–869. https://doi.org/10.1046/j.1365-2958.1996.521416.x.

Håvarstein, L.S., Martin, B., Johnsborg, O., Granadel, C., and Claverys, J.-P. (2006). New insights into the pneumococcal fratricide: relationship to clumping and identification of a novel immunity factor. Mol. Microbiol. 59, 1297–1307. https://doi.org/10.1111/j.1365-2958.2005.05021.x.

Heß, N., Waldow, F., Kohler, T.P., Rohde, M., Kreikemeyer, B., Gómez-Mejia, A., Hain, T., Schwudke, D., Vollmer, W., Hammerschmidt, S., et al. (2017). Lipoteichoic acid deficiency permits normal growth but impairs virulence of Streptococcus pneumoniae. Nat. Commun. 8, 2093. https://doi.org/10.1038/s41467-017-01720-z.

Hui, F.M., and Morrison, D.A. (1991). Genetic transformation in Streptococcus pneumoniae: nucleotide sequence analysis shows comA, a gene required for competence induction, to be a member of the bacterial ATP-dependent transport protein family. J. Bacteriol. 173, 372–381. https://doi.org/10.1128/jb.173.1.372-381.1991.

Iovino, F., Engelen-Lee, J.-Y., Brouwer, M., van de Beek, D., van der Ende, A., Valls Seron, M., Mellroth, P., Muschiol, S., Bergstrand, J., Widengren, J., et al. (2017). pIgR and PECAM-1 bind to pneumococcal adhesins RrgA and PspC mediating bacterial brain invasion. J. Exp. Med. 214, 1619–1630. https://doi.org/10.1084/jem.20161668.

Jim, K.K., Aprianto, R., Domenech, A., Kurushima, J., Beek, D. van de, Vandenbroucke-Grauls, C.M.J.E., Bitter, W., and Veening, J.-W. (2022). Pneumolysin promotes host cell necroptosis and bacterial competence during pneumococcal meningitis as shown by whole animal dual RNA-seq. 2022.02.10.479878. https://doi.org/10.1101/2022.02.10.479878.

Johnsborg, O., and Håvarstein, L.S. (2009). Pneumococcal LytR, a protein from the LytR-CpsA-Psr family, is essential for normal septum formation in Streptococcus pneumoniae. J. Bacteriol. 191, 5859–5864. https://doi.org/10.1128/JB.00724-09.

Johnsborg, O., Eldholm, V., Bjørnstad, M.L., and Håvarstein, L.S. (2008). A predatory mechanism dramatically increases the efficiency of lateral gene transfer in Streptococcus pneumoniae and related commensal species. Mol. Microbiol. 69, 245–253. https://doi.org/10.1111/j.1365-2958.2008.06288.x.

Johnston, C., Martin, B., Fichant, G., Polard, P., and Claverys, J.-P. (2014). Bacterial transformation: distribution, shared mechanisms and divergent control. Nat. Rev. Microbiol. 12, 181–196. https://doi.org/10.1038/nrmicro3199.

de Jong, I.G., Beilharz, K., Kuipers, O.P., and Veening, J.-W. (2011). Live Cell Imaging of Bacillus subtilis and Streptococcus pneumoniae using Automated Time-lapse Microscopy. J. Vis. Exp. JoVE 3145. https://doi.org/10.3791/3145.

Junges, R., Maienschein-Cline, M., Morrison, D.A., and Petersen, F.C. (2019). Complete Genome Sequence of Streptococcus pneumoniae Serotype 19F Strain EF3030. Microbiol. Resour. Announc. 8, e00198–19. https://doi.org/10.1128/MRA.00198-19.

Kadioglu, A., Weiser, J.N., Paton, J.C., and Andrew, P.W. (2008). The role of Streptococcus pneumoniae virulence factors in host respiratory colonization and disease. Nat. Rev. Microbiol. 6, 288–301. https://doi.org/10.1038/nrmicro1871.

Kausmally, L., Johnsborg, O., Lunde, M., Knutsen, E., and Håvarstein, L.S. (2005). Choline-binding protein D (CbpD) in Streptococcus pneumoniae is essential for competence-induced cell lysis. J. Bacteriol. 187, 4338–4345. https://doi.org/10.1128/JB.187.13.4338-4345.2005.

Kawai, Y., Marles-Wright, J., Cleverley, R.M., Emmins, R., Ishikawa, S., Kuwano, M., Heinz, N., Bui, N.K., Hoyland, C.N., Ogasawara, N., et al. (2011). A widespread family of bacterial cell wall assembly proteins. EMBO J. 30, 4931–4941. https://doi.org/10.1038/emboj.2011.358.

Keller, L.E., Rueff, A.-S., Kurushima, J., and Veening, J.-W. (2019). Three New Integration Vectors and Fluorescent Proteins for Use in the Opportunistic Human Pathogen Streptococcus pneumoniae. Genes 10. https://doi.org/10.3390/genes10050394.

Kjos, M., Miller, E., Slager, J., Lake, F.B., Gericke, O., Roberts, I.S., Rozen, D.E., and Veening, J.-W. (2016). Expression of Streptococcus pneumoniae Bacteriocins Is Induced by Antibiotics via Regulatory Interplay with the Competence System. PLoS Pathog. 12, e1005422. https://doi.org/10.1371/journal.ppat.1005422.

Larson, T.R., and Yother, J. (2017). Streptococcus pneumoniae capsular polysaccharide is linked to peptidoglycan via a direct glycosidic bond to β-D-N-acetylglucosamine. Proc. Natl. Acad. Sci. U. S. A. 114, 5695–5700. https://doi.org/10.1073/pnas.1620431114.

Lee, M.S., and Morrison, D.A. (1999). Identification of a new regulator in Streptococcus pneumoniae linking quorum sensing to competence for genetic transformation. J. Bacteriol. 181, 5004–5016. https://doi.org/10.1128/JB.181.16.5004-5016.1999.

Lin, J., Park, P., Li, H., Oh, M.W., Dobrucki, I.T., Dobrucki, W., and Lau, G.W. (2020). Streptococcus pneumoniae Elaborates Persistent and Prolonged Competent State during Pneumonia-Derived Sepsis. Infect. Immun. 88, e00919–19. https://doi.org/10.1128/IAI.00919-19.

Liu, X., Gallay, C., Kjos, M., Domenech, A., Slager, J., van Kessel, S.P., Knoops, K., Sorg, R.A., Zhang, J.-R., and Veening, J.-W. (2017). High-throughput CRISPRi phenotyping identifies new essential genes in Streptococcus pneumoniae. Mol. Syst. Biol. 13, 931. https://doi.org/10.15252/msb.20167449.

Liu, X., Kimmey, J.M., Matarazzo, L., de Bakker, V., Van Maele, L., Sirard, J.-C., Nizet, V., and Veening, J.-W. (2021). Exploration of Bacterial Bottlenecks and Streptococcus pneumoniae Pathogenesis by CRISPRi-Seq. Cell Host Microbe 29, 107–120.e6. https://doi.org/10.1016/j.chom.2020.10.001.

Mann, B., Orihuela, C., Antikainen, J., Gao, G., Sublett, J., Korhonen, T.K., and Tuomanen, E. (2006). Multifunctional role of choline binding protein G in pneumococcal pathogenesis. Infect. Immun. 74, 821–829. https://doi.org/10.1128/IAI.74.2.821-829.2006.

Martin, B., Soulet, A.-L., Mirouze, N., Prudhomme, M., Mortier-Barrière, I., Granadel, C., Noirot-Gros, M.-F., Noirot, P., Polard, P., and Claverys, J.-P. (2013). ComE/ComE~P interplay dictates activation or extinction status of pneumococcal X-state (competence). Mol. Microbiol. 87, 394–411. https://doi.org/10.1111/mmi.12104.

Matarazzo, L., Casilag, F., Porte, R., Wallet, F., Cayet, D., Faveeuw, C., Carnoy, C., and Sirard, J.-C. (2019). Therapeutic Synergy Between Antibiotics and Pulmonary Toll-Like Receptor 5 Stimulation in Antibiotic-Sensitive or -Resistant Pneumonia. Front. Immunol. 10, 723. https://doi.org/10.3389/fimmu.2019.00723.

Park, S.-S., Gonzalez-Juarbe, N., Martínez, E., Hale, J.Y., Lin, Y.-H., Huffines, J.T., Kruckow, K.L., Briles, D.E., and Orihuela, C.J. (2021). Streptococcus pneumoniae Binds to Host Lactate Dehydrogenase via PspA and PspC To Enhance Virulence. MBio 12, e00673–21. https://doi.org/10.1128/mBio.00673-21.

Pestova, E.V., Håvarstein, L.S., and Morrison, D.A. (1996). Regulation of competence for genetic transformation in Streptococcus pneumoniae by an auto-induced peptide pheromone and a two-component regulatory system. Mol. Microbiol. 21, 853–862. https://doi.org/10.1046/j.1365-2958.1996.501417.x.

Peterson, S.N., Sung, C.K., Cline, R., Desai, B.V., Snesrud, E.C., Luo, P., Walling, J., Li, H., Mintz, M., Tsegaye, G., et al. (2004). Identification of competence pheromone responsive genes in Streptococcus pneumoniae by use of DNA microarrays. Mol. Microbiol. 51, 1051–1070. https://doi.org/10.1046/j.1365-2958.2003.03907.x.

Rosenow, C., Ryan, P., Weiser, J.N., Johnson, S., Fontan, P., Ortqvist, A., and Masure, H.R. (1997). Contribution of novel choline-binding proteins to adherence, colonization and immunogenicity of Streptococcus pneumoniae. Mol. Microbiol. 25, 819–829. https://doi.org/10.1111/j.1365-2958.1997.mmi494.x.

Schmidt, F., Kakar, N., Meyer, T.C., Depke, M., Masouris, I., Burchhardt, G., Gómez-Mejia, A., Dhople, V., Håvarstein, L.S., Sun, Z., et al. (2019). In vivo proteomics identifies the competence regulon and AliB oligopeptide transporter as pathogenic factors in pneumococcal meningitis. PLoS Pathog. 15, e1007987. https://doi.org/10.1371/journal.ppat.1007987.

Selinger, D.S., and Reed, W.P. (1979). Pneumococcal adherence to human epithelial cells. Infect. Immun. 23, 545–548. https://doi.org/10.1128/iai.23.2.545-548.1979.

Seo, H.S., Cartee, R.T., Pritchard, D.G., and Nahm, M.H. (2008). A new model of pneumococcal lipoteichoic acid structure resolves biochemical, biosynthetic, and serologic inconsistencies of the current model. J. Bacteriol. 190, 2379–2387. https://doi.org/10.1128/JB.01795-07.

Slager, J., Kjos, M., Attaiech, L., and Veening, J.-W. (2014). Antibiotic-induced replication stress triggers bacterial competence by increasing gene dosage near the origin. Cell 157, 395–406. https://doi.org/10.1016/j.cell.2014.01.068.

Slager, J., Aprianto, R., and Veening, J.-W. (2018). Deep genome annotation of the opportunistic human pathogen Streptococcus pneumoniae D39. Nucleic Acids Res. 46, 9971–9989. https://doi.org/10.1093/nar/gky725.

Slager, J., Aprianto, R., and Veening, J.-W. (2019). Refining the Pneumococcal Competence Regulon by RNA Sequencing. J. Bacteriol. 201. https://doi.org/10.1128/JB.00780-18.

Stefanović, C., Hager, F.F., and Schäffer, C. (2021). LytR-CpsA-Psr Glycopolymer Transferases: Essential Bricks in Gram-Positive Bacterial Cell Wall Assembly. Int. J. Mol. Sci. 22, E908. https://doi.org/10.3390/ijms22020908.

Steinmoen, H., Knutsen, E., and Håvarstein, L.S. (2002). Induction of natural competence in Streptococcus pneumoniae triggers lysis and DNA release from a subfraction of the cell population. Proc. Natl. Acad. Sci. U. S. A. 99, 7681–7686. https://doi.org/10.1073/pnas.112464599.

Straume, D., Stamsås, G.A., Salehian, Z., and Håvarstein, L.S. (2017a). Overexpression of the fratricide immunity protein ComM leads to growth inhibition and morphological abnormalities in Streptococcus pneumoniae. Microbiol. Read. Engl. 163, 9–21. https://doi.org/10.1099/mic.0.000402.

Straume, D., Stamsås, G.A., Berg, K.H., Salehian, Z., and Håvarstein, L.S. (2017b). Identification of pneumococcal proteins that are functionally linked to penicillin-binding protein 2b (PBP2b). Mol. Microbiol. 103, 99–116. https://doi.org/10.1111/mmi.13543.

Straume, D., Piechowiak, K.W., Olsen, S., Stamsås, G.A., Berg, K.H., Kjos, M., Heggenhougen, M.V., Alcorlo, M., Hermoso, J.A., and Håvarstein, L.S. (2020). Class A PBPs have a distinct and unique role in the construction of the pneumococcal cell wall. Proc. Natl. Acad. Sci. U. S. A. 117, 6129–6138. https://doi.org/10.1073/pnas.1917820117.

Subramanian, K., Henriques-Normark, B., and Normark, S. (2019). Emerging concepts in the pathogenesis of the Streptococcus pneumoniae: From nasopharyngeal colonizer to intracellular pathogen. Cell. Microbiol. 21, e13077. https://doi.org/10.1111/cmi.13077.

Swiatlo, E., Champlin, F.R., Holman, S.C., Wilson, W.W., and Watt, J.M. (2002). Contribution of Choline-Binding Proteins to Cell Surface Properties of Streptococcus pneumoniae. Infect. Immun. 70, 412–415. https://doi.org/10.1128/IAI.70.1.412-415.2002.

Tomasz, A. (1967). Choline in the cell wall of a bacterium: novel type of polymer-linked choline in Pneumococcus. Science 157, 694–697. https://doi.org/10.1126/science.157.3789.694.

Veening, J.-W., and Blokesch, M. (2017). Interbacterial predation as a strategy for DNA acquisition in naturally competent bacteria. Nat. Rev. Microbiol. 15, 621–629. https://doi.org/10.1038/nrmicro.2017.66.

Wahl, B., O’Brien, K.L., Greenbaum, A., Majumder, A., Liu, L., Chu, Y., Lukšić, I., Nair, H., McAllister, D.A., Campbell, H., et al. (2018). Burden of Streptococcus pneumoniae and Haemophilus influenzae type b disease in children in the era of conjugate vaccines: global, regional, and national estimates for 2000-15. Lancet Glob. Health 6, e744–e757. https://doi.org/10.1016/S2214-109X(18)30247-X.

Weidenmaier, C., and Peschel, A. (2008). Teichoic acids and related cell-wall glycopolymers in Gram-positive physiology and host interactions. Nat. Rev. Microbiol. 6, 276–287. https://doi.org/10.1038/nrmicro1861.

Weiser, J.N., Ferreira, D.M., and Paton, J.C. (2018). Streptococcus pneumoniae: transmission, colonization and invasion. Nat. Rev. Microbiol. 16, 355–367. https://doi.org/10.1038/s41579-018-0001-8.

Wholey, W.-Y., Kochan, T.J., Storck, D.N., and Dawid, S. (2016). Coordinated Bacteriocin Expression and Competence in Streptococcus pneumoniae Contributes to Genetic Adaptation through Neighbor Predation. PLoS Pathog. 12, e1005413. https://doi.org/10.1371/journal.ppat.1005413.

Wyres, K.L., Lambertsen, L.M., Croucher, N.J., McGee, L., von Gottberg, A., Liñares, J., Jacobs, M.R., Kristinsson, K.G., Beall, B.W., Klugman, K.P., et al. (2013). Pneumococcal capsular switching: a historical perspective. J. Infect. Dis. 207, 439–449. https://doi.org/10.1093/infdis/jis703.

Ye, W., Zhang, J., Shu, Z., Yin, Y., Zhang, X., and Wu, K. (2018). Pneumococcal LytR Protein Is Required for the Surface Attachment of Both Capsular Polysaccharide and Teichoic Acids: Essential for Pneumococcal Virulence. Front. Microbiol. 9, 1199. https://doi.org/10.3389/fmicb.2018.01199.

Zhang, J.-R., Mostov, K.E., Lamm, M.E., Nanno, M., Shimida, S., Ohwaki, M., and Tuomanen, E. (2000). The Polymeric Immunoglobulin Receptor Translocates Pneumococci across Human Nasopharyngeal Epithelial Cells. Cell 102, 827–837. https://doi.org/10.1016/S0092-8674(00)00071-4.

