## Supplemental Figures S1-S8 for "Competence remodels the pneumococcal cell wall providing resistance to fratricide and surface exposing key virulence factors"

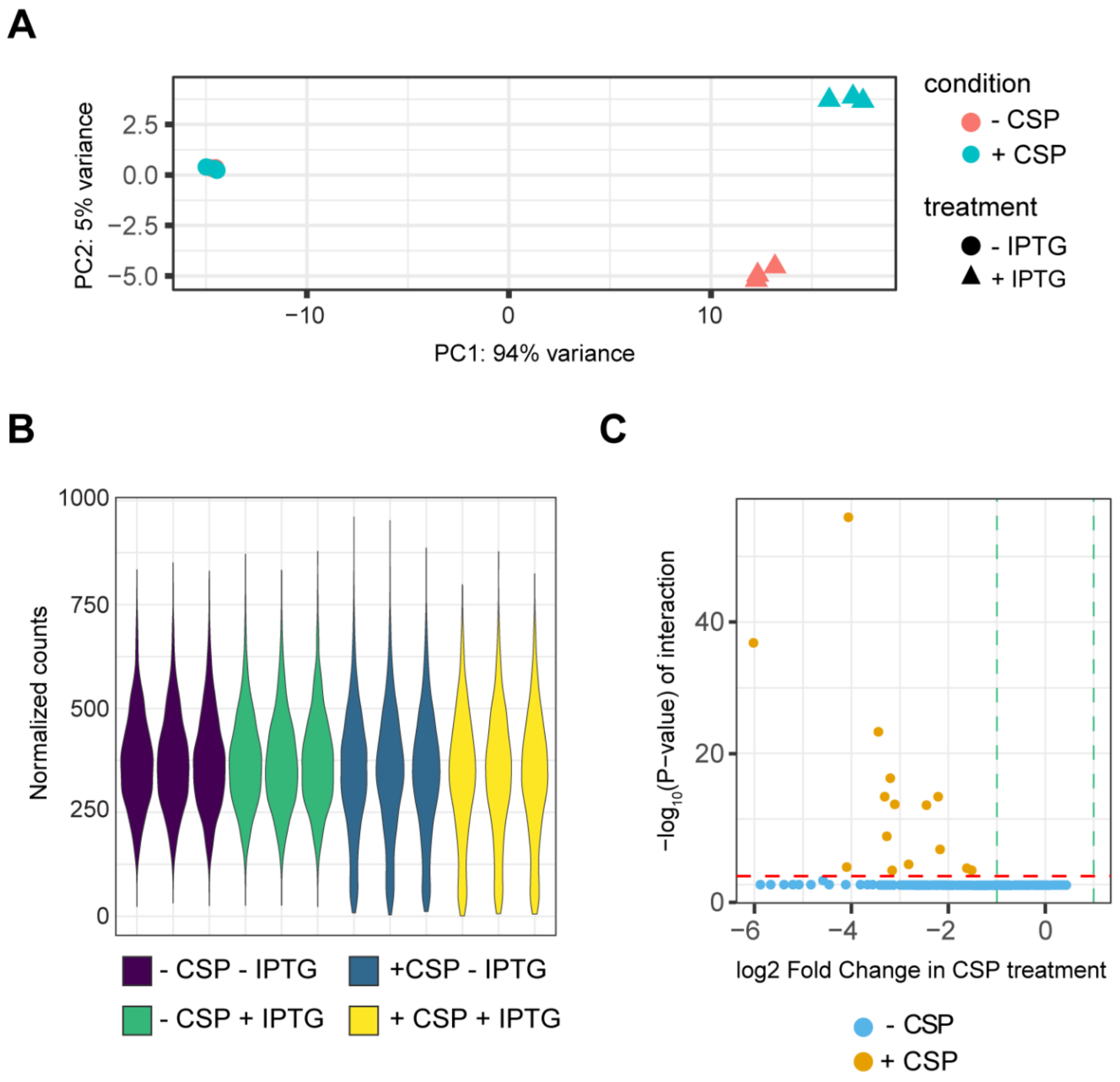


**Fig. S1. Evaluation of fitness cost during competence using CRISPRi pool screen. A-B**) Variance of the replicates. **C**) IPTG main effect in competence by interaction of p−values. Tested against fold change of 2 (green lines) with alpha of 0.05 (red line).

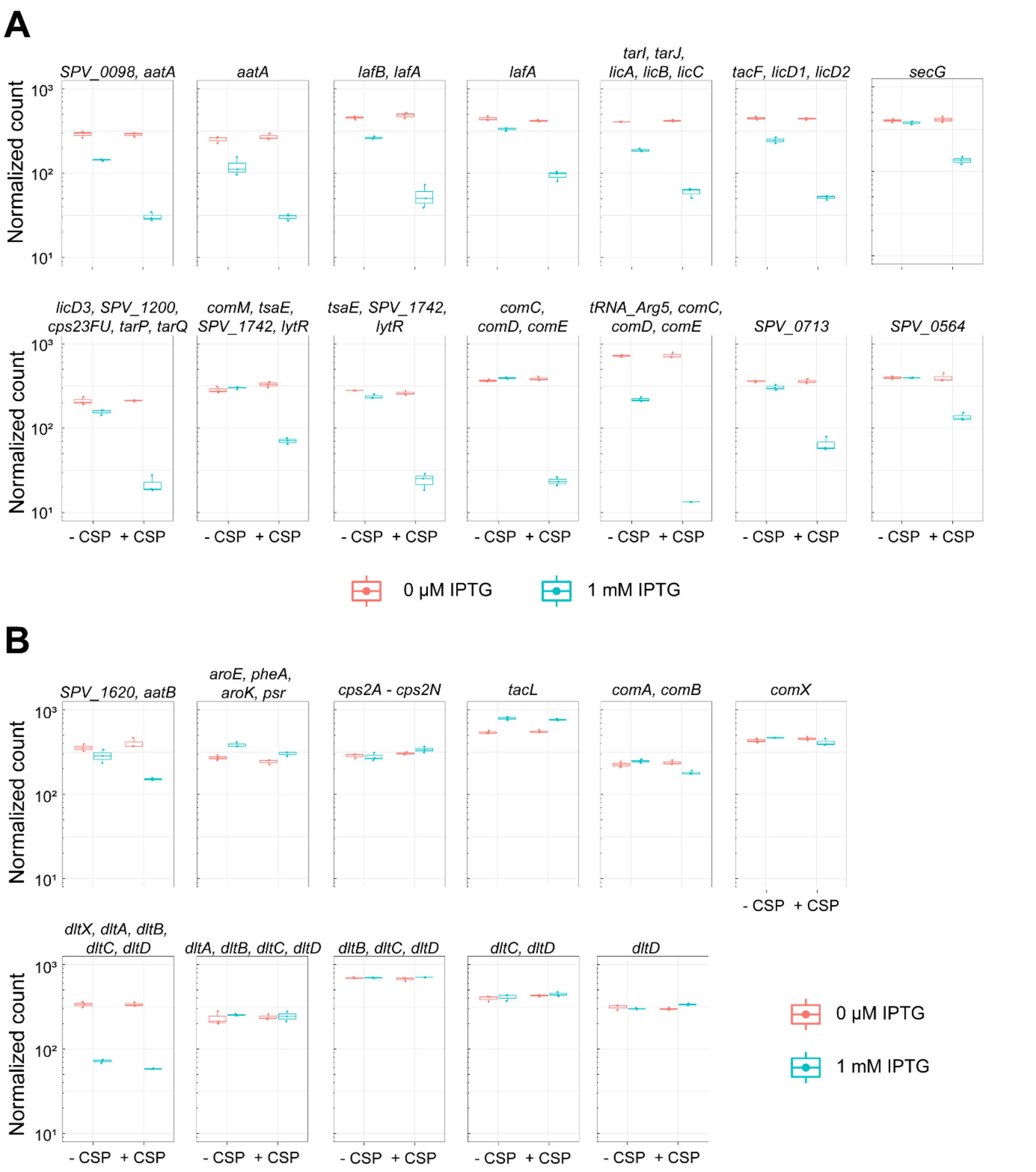


**Figure S2. Normalized counts of sgRNAs related to competence and teichoic acid (TA) synthesis. A)** sgRNAs with a significant fitness cost during competence**. B)** other sgRNAs related to competence or TA synthesis with no fitness cost. Fitness cost was evaluated as described before (de Bakker et al., 2022) (see methods for more details).

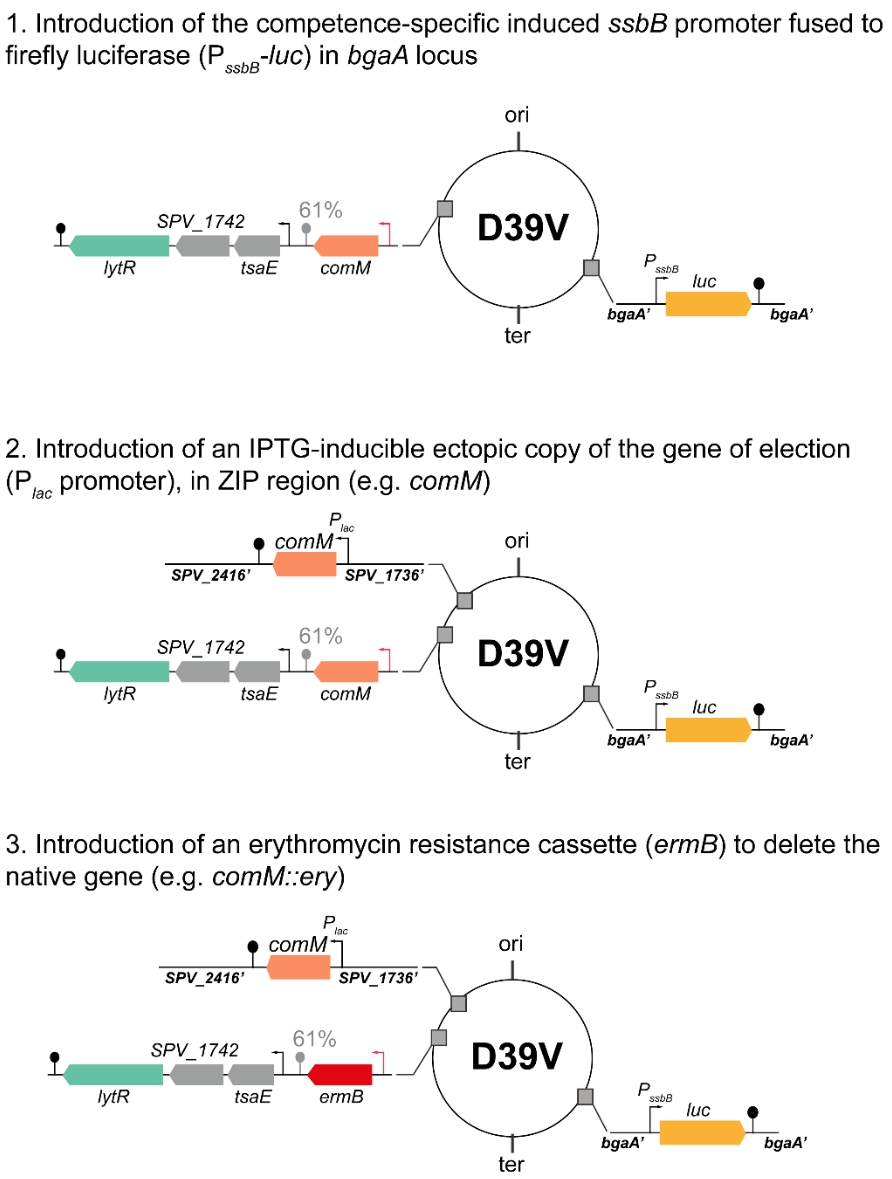


**Figure S3.** Design of inducible systems.

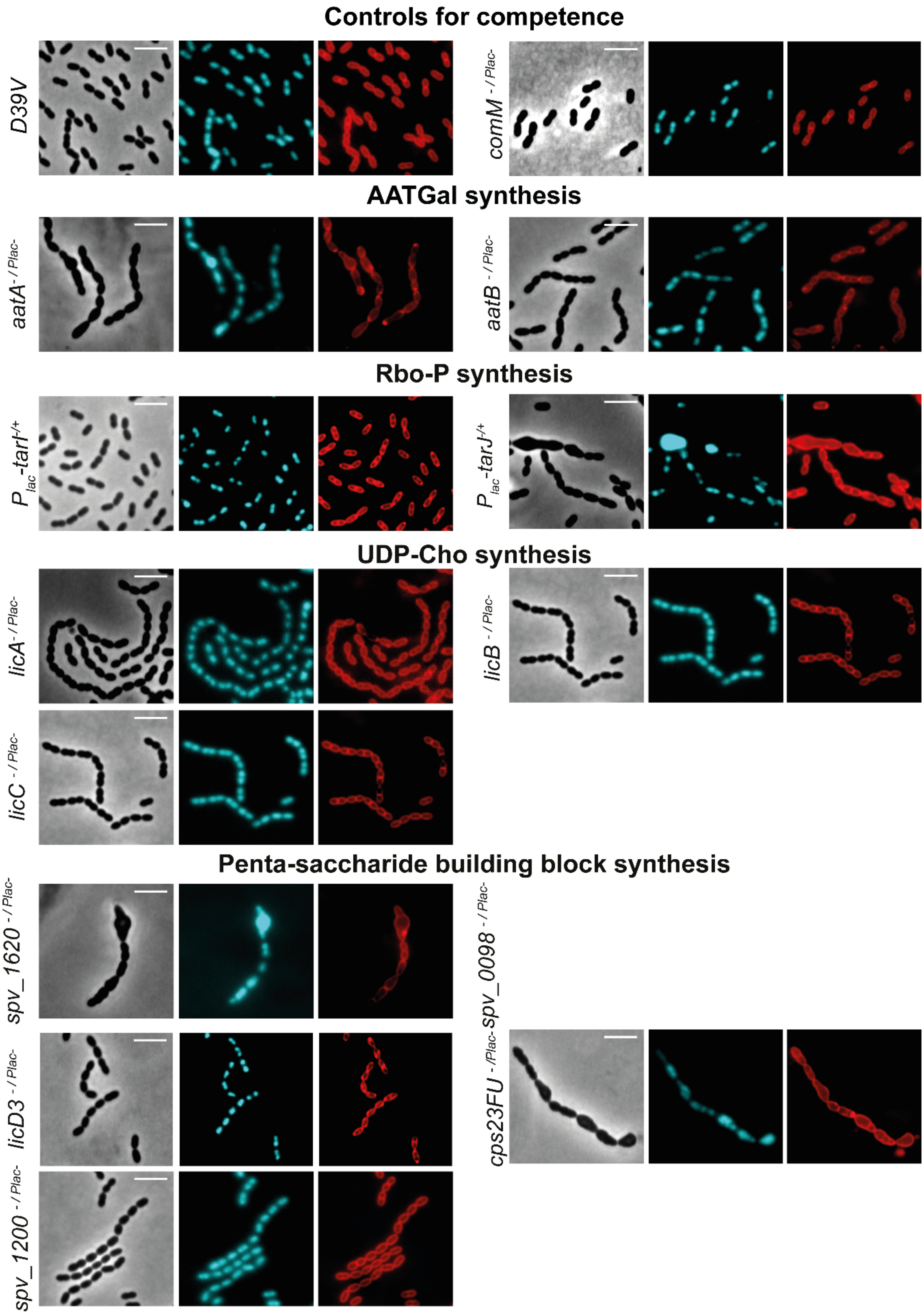

Controls for competence
D39V
comM -/ Plac-
AATGal synthesis
aatA -/ Plac-
aatB -/ Plac-
Rbo-P synthesis
Plac-tarI-/+
Plac-tarJ-/+
UDP-Cho synthesis
licA -/ Plac-
licB -/ Plac-
licC -/ Plac-
Penta-saccharide building block synthesis
spv_1620 -/ Plac-
spv_0098 -/ Plac-
licD3 -/ Plac-
cps23FU -/Plac-
spv_1200 -/ Plac-

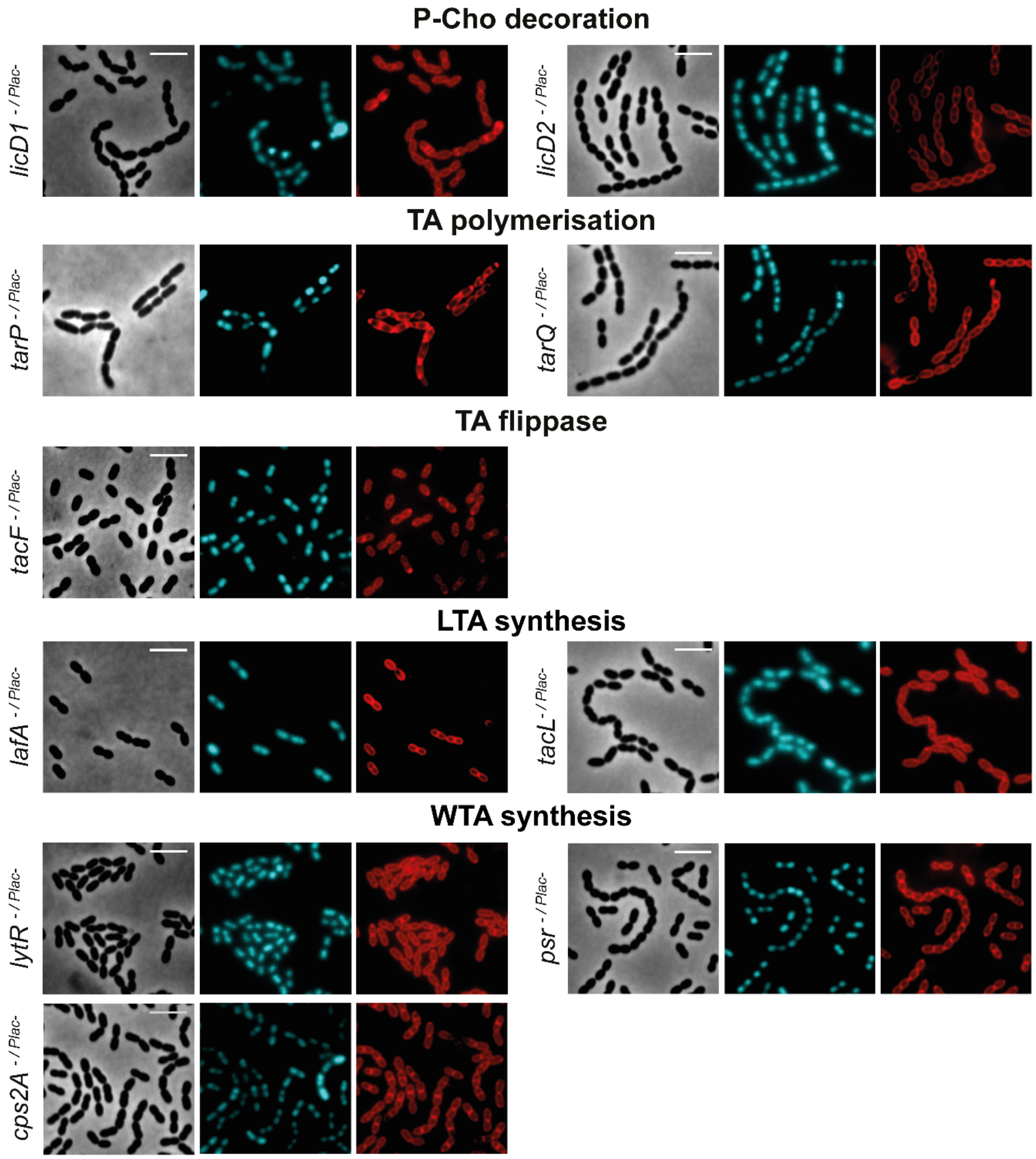


**Figure S4.** Morphological changes were examined with fluorescence microscopy, and representative micrographs are shown. Phase contrast, DAPI staining, and Nile red staining are displayed.

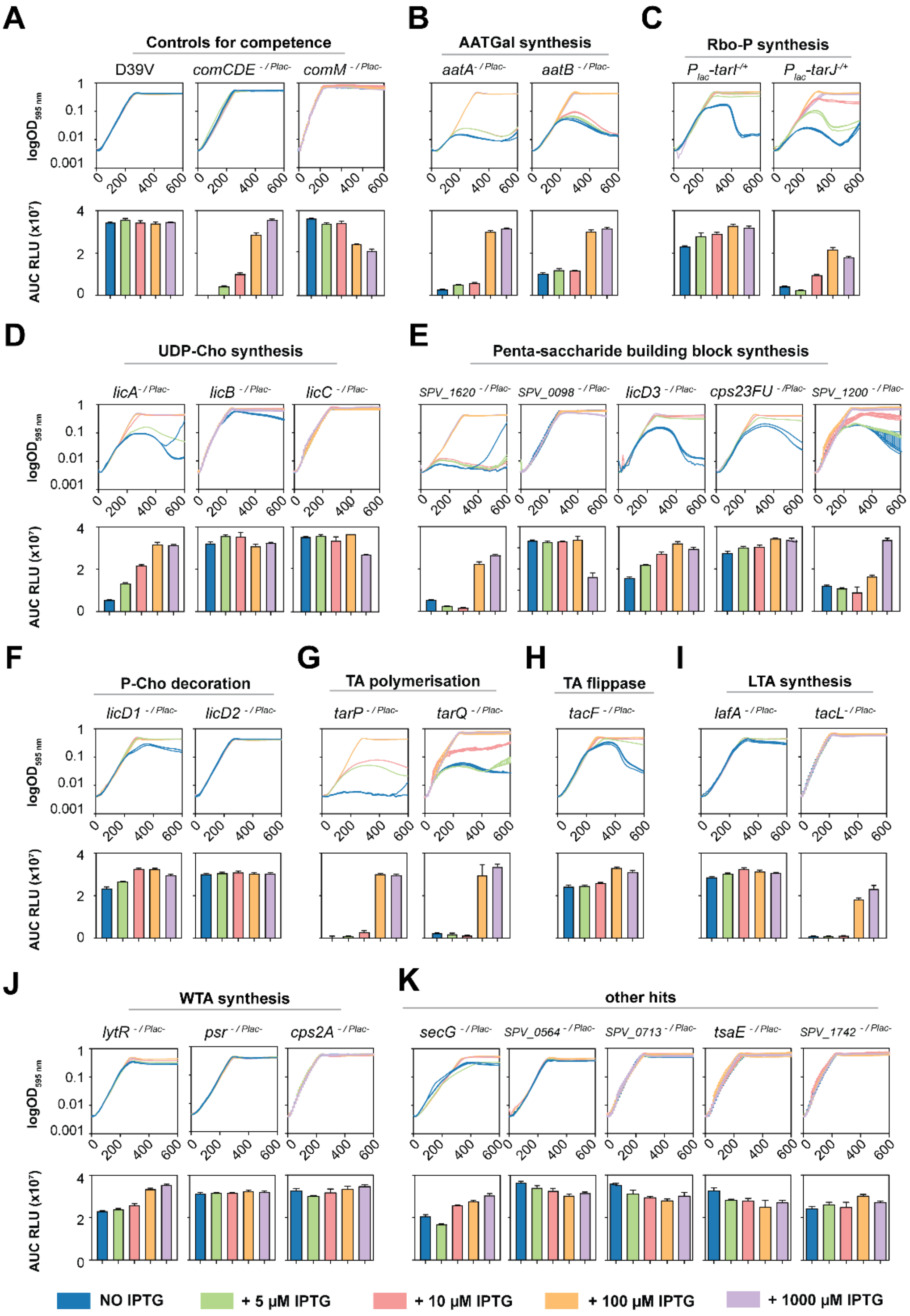


**Figure S5. Natural competence development in competence and teichoic acid related genes.** Every strain contains a depletion system by ectopically expressing the indicated gene under control of the Plac IPTG-inducible promoter, and the deletion of the gene from its native location. Top, growth curves in absence of the protein (blue = No IPTG) or with different IPTG concentrations. Bottom, area under the curve (AUC) of relative luminescence values (RLU).

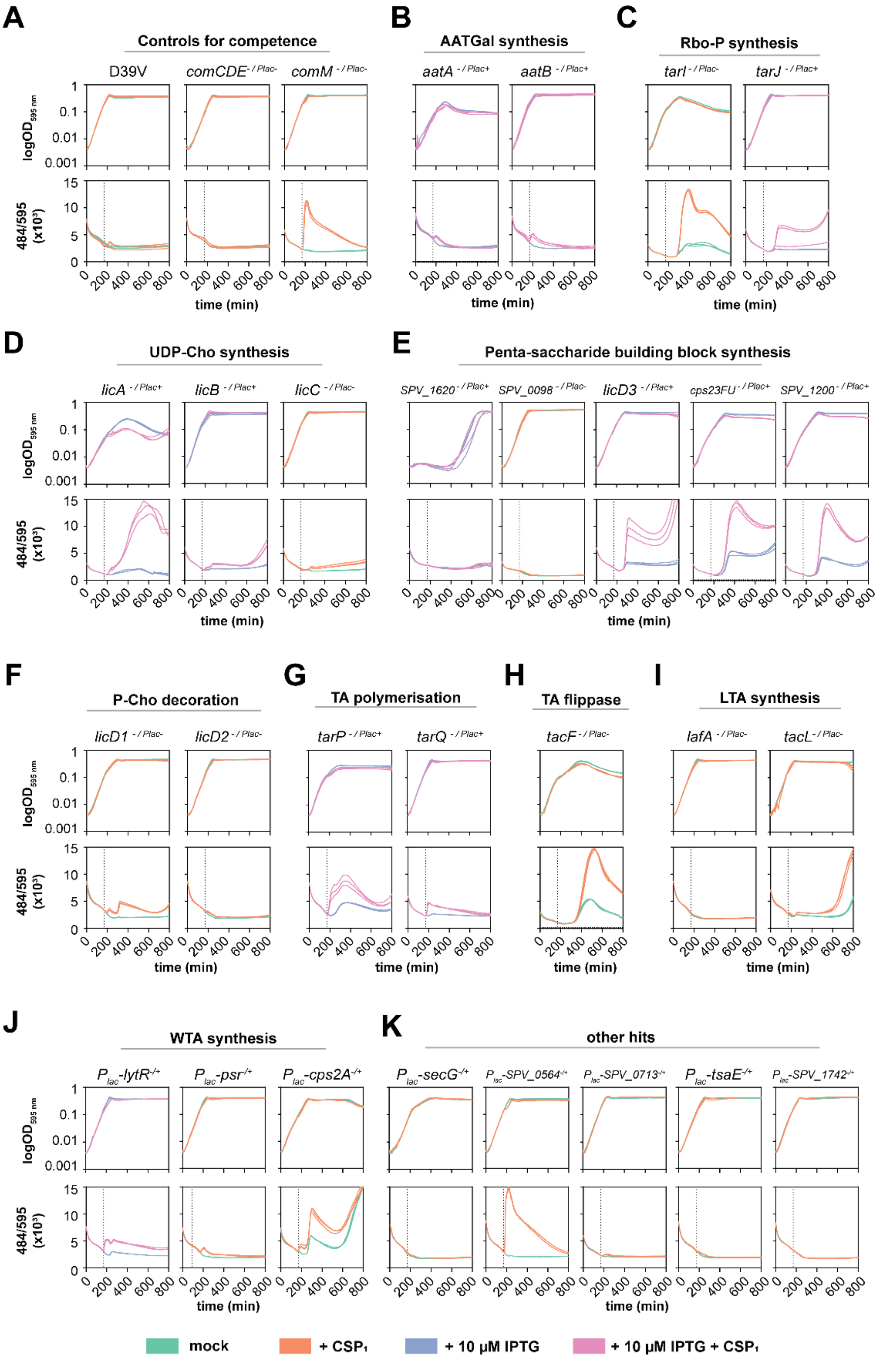


**Figure S6. Detection of cell lysis.** Individual strains were grown in C+Y pH 6.8 to avoid natural competence development in presence of SYTOX™ Green Dead Cell Stain dye. When $OD_{595\ nm}$ reached ~ 0.1, 100 ng/ml of $CSP_1$ was added to induce competence. For those genes showing less cell lysis in our setup, experiment was performed in absence of IPTG (green and orange). For those essential with severe growth defect, 10 µM IPTG was added to maintain a mild expression of the protein and avoid cell lysis in absence of CSP due to the essentiality of the gene (blue and pink).

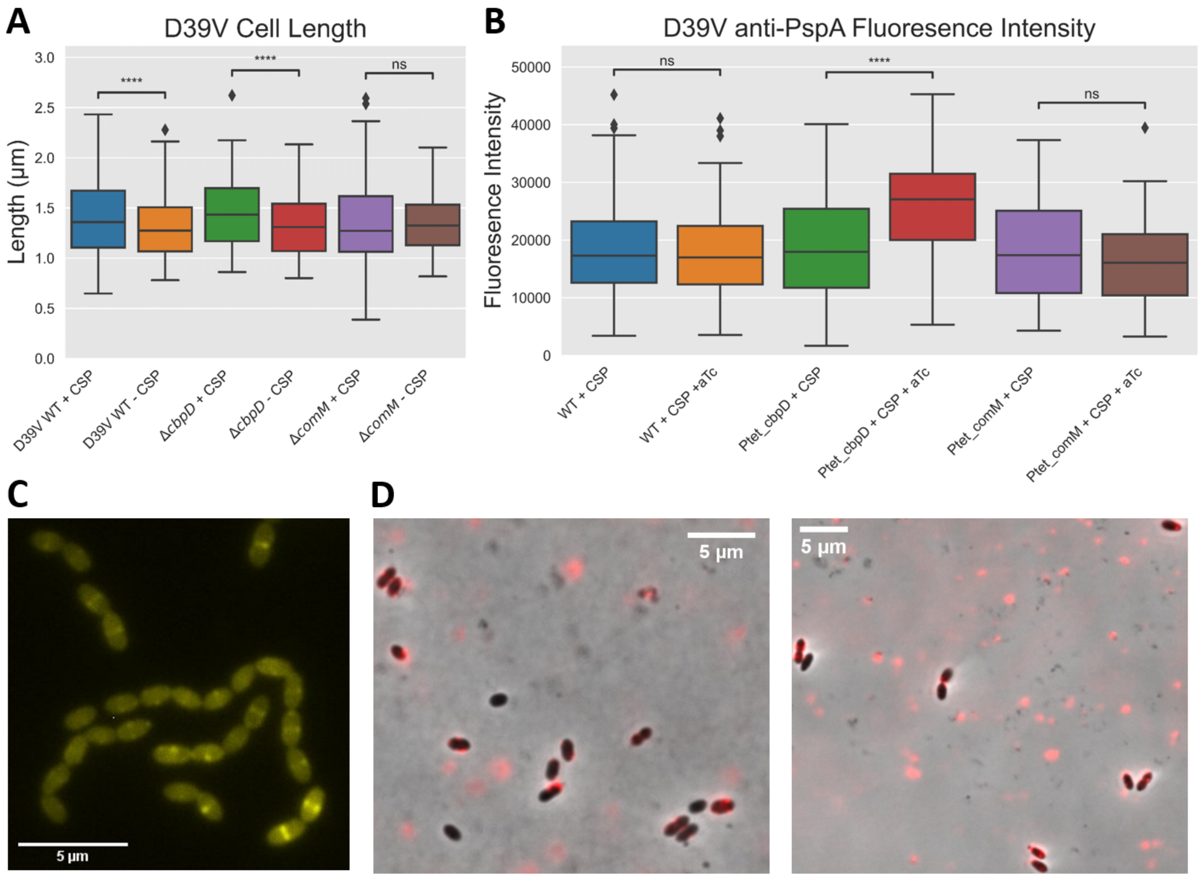


**Figure S7. Cell length measurements and PspA overexpression immunofluorescence after competence induction, PspA immunofluorescence of a Δ*cps* mutant. A**) D39V cells were grown to OD 0.1 in C+Y medium pH 7.4, then exposed to 100 ng/ml or 0 ng/ml of CSP-1 for 30 min. Samples were subjected to epifluorescence microscopy, where phase contrast images were used to measure cell length. Diamond symbols represent outlier individual cell length measurements. Asterisks show statistically significant differences in cell length (Mann-Whitney U test) (see methods section for more details). **B**) D39V strains containing aTc inducible promoters coupled to *cbpD* or *comM* were grown to OD 0.1 in C+Y medium pH 7.4, then exposed to 100 ng/ml or 0 ng/ml of CSP-1 for 30 min and/or 100ng/ml aTc**.** Cells were then stained with primary antibodies raised against PspA and then Goat anti-rabbit IgG (H+L) Alexa 555, both at 1/500 dilutions and subjected to epifluorescence microscopy. Fluorescence intensity based on phase-contrast and fluorescence composite images was measured. Diamond symbols represent outlier individual fluorescence measurements. Asterisks show statistically significant differences in cell length (Mann-Whitney U test) (see methods section for more details). **C**) Representative phase contrast and fluorescence composite image taken of D39V strain with *comM* genetically tagged with YFP at its C-terminus. **D**) D39V Δ*cps* cells were grown to OD 0.1 in C+Y medium pH 7.4, then exposed to 100 ng/ml (left image) or 0 ng/ml (right image) of CSP-1 for 30 min. Cells were then stained with primary antibodies raised against PspA, and then Goat anti-rabbit IgG (H+L) Alexa 555, both at 1/500 dilutions and subjected to epifluorescence microscopy (see methods section for more details).

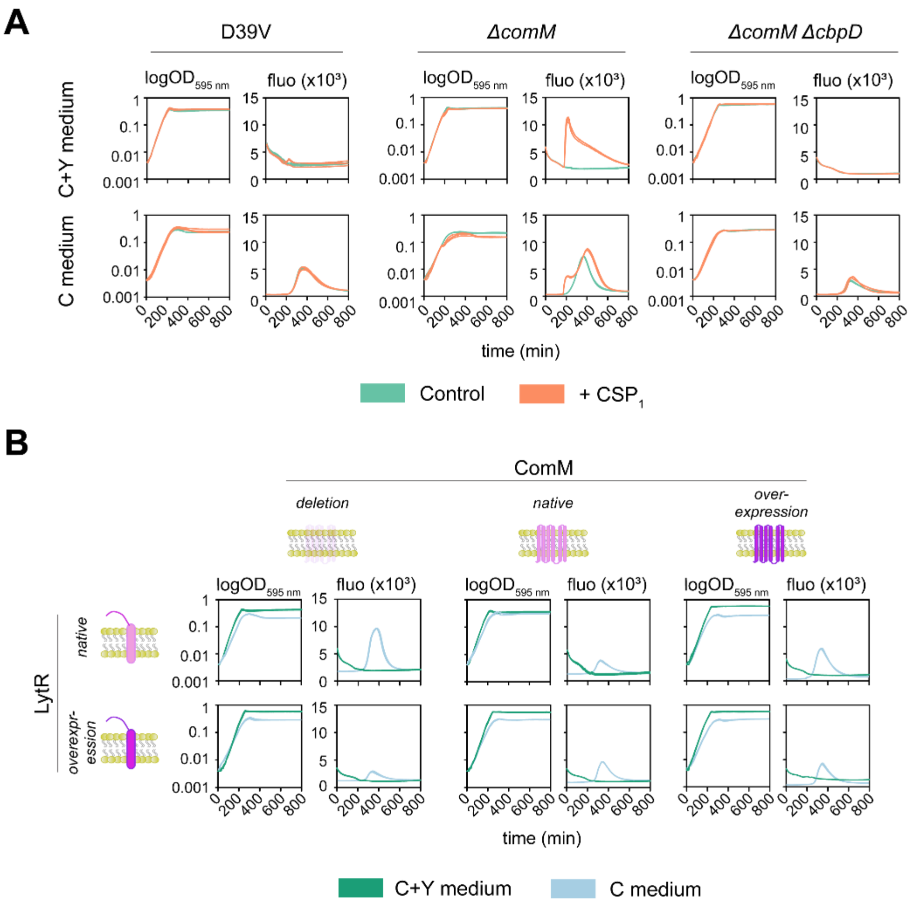


**Figure S8. Cell lysis detection in C+Y and C media. A)** Cell lysis detection in D39V, Δ*comM* and double Δ*comM*Δ*cbpD* strains. Individual strains were grown in C+Y medium (top) or C medium (bottom) at pH 6.8 to avoid natural competence development in presence of SYTOX™ Green Dead Cell Stain dye. When cell cultures reached $OD_{595\ nm}$ 0.1 (~ after 170 min), 100 ng/ml of $CSP_1$ was added to induce competence (orange lines). Three biological replicates per condition are shown. **B)** Evaluation of cell lysis in the strains used for the radioactive assay (Figure 4). Cells were grown in the indicated medium in presence of SYTOX™ Green Dead Cell Stain dye. Three biological replicates per condition are shown.
